# Base-modified nucleotides mediate immune signaling in bacteria

**DOI:** 10.1101/2024.12.20.629578

**Authors:** Zhifeng Zeng, Zeyu Hu, Ruiliang Zhao, Jikai Rao, Mario Rodríguez Mestre, Yanqiu Liu, Shunhang Liu, Hao Feng, Yu Chen, Huan He, Nuo Chen, Jinshui Zheng, Donghai Peng, Min Luo, Qunxin She, Rafael Pinilla-Redondo, Wenyuan Han

## Abstract

Signaling from pathogen sensing to effector activation is a fundamental principle of cellular immunity. While cyclic (oligo)nucleotides have emerged as key signaling molecules, the existence of other messengers remains largely unexplored. Here, we reveal a bacterial anti-phage system that mediates immune signaling through nucleobase modification. Immunity is triggered by phage nucleotide kinases, which, combined with the system-encoded adenosine deaminase, produce deoxyinosine 5’-triphosphate (dITP) as immune messengers. The dITP signal activates downstream effector to mediate cellular NAD^+^ depletion, resulting in population-level defense through the death of infected cells. To counteract immune signaling, phages deploy specialized enzymes that deplete cellular dAMP, the precursor of dITP messengers. Our findings uncover a nucleobase modification-based anti-phage signaling pathway, establishing noncanonical nucleotides as a new type of immune messengers in bacteria.

---

All cellular life faces constant threats from pathogen invasion, including viruses, driving the evolution of sophisticated immune systems (*1, 2*). Antiviral systems typically function by relaying pathogen detection to immune effector activation. This signal transduction can occur through mechanisms such as conformational changes or physical interaction of immune components, and the synthesis of signaling molecules (*2, 3*). Diverse immune pathways have converged on using signaling molecule synthesis due to its advantages in amplifying minimal infection cues and allowing rapid system diversification through sensor and effector swapping (*4-6*). Indeed, immune signaling is widespread across prokaryotic and eukaryotic antiviral systems, including the cGAS-STING, OAS-RNaseL, and Toll/interleukin-1 receptor (TIR)-dependent systems in animals and plants (*5, 7*), as well as the type III CRISPR-Cas (*8, 9*), CBASS (*10, 11*), Thoeris (*12*), and Pycsar (*13*) systems in bacteria and archaea. These systems share conceptually similar signal transduction strategies, culminating in the activation of downstream cell death or dormancy effectors that function through diverse mechanisms, including degradation of cellular nucleic acids (*8, 9*), disruption of membrane integrity (*10*) and depletion of the essential metabolic cofactor NAD^+^ (*12*). Remarkably, the involved immune signaling molecules are conserved from prokaryotes to animals and plants, typically including cyclic-(oligo)nucleotides and cyclic adenine diphosphate ribose (cADPR) variants. The cADPR variants are generated via processing of NAD^+^ by TIR domains (*7, 12*), while nucleotide transferases and cyclases catalyze the synthesis of cyclic- (oligo)nucleotides (*5, 8-11, 13*). However, the existence of other types of signaling molecules remains unclear.

In this study, we discover and functionally characterize a three-gene bacterial anti-phage defense system. The system specifically produces deoxyinosine 5’-triphosphate (dITP) as an immune signal messenger that activates an immune effector complex to deplete cellular NAD^+^, leading to population-level protection through the suicide of infected cells. Significantly different from all known immune signaling systems, the three-gene system hijacks phage nucleotide kinases to complete a cascade reaction for signaling molecule synthesis. Inspired by this feature, we name the three-gene system after Kongming, a great military strategist known in a story “borrowing arrows from enemy”, and designate the three genes *komA*, *komB*, *komC* respectively. Finally, we also identify a phage-encoded anti-defense protein that subverts dITP signaling by degrading dAMP, the precursor for dITP synthesis. Our findings establish a novel type of immune signaling in bacteria and ascribe an unprecedented biological function to the noncanonical nucleotide dITP.

## Results

### Identification of a three-gene anti-phage system

Defense systems have a tendency to cluster in bacterial genomes within so-called “defense islands” (*14*), which provides a powerful strategy to identify new defense systems (*15-17*). Through comparative genomics analyses, we identified a three-gene operon that is sometimes located in the neighborhood of characterized defense systems (Fig. 1A). This three-gene system (hereafter Kongming) encodes a predicted adenosine deaminase (pfam PF00962, KomA), a HAM1-like non-canonical purine NTP pyrophosphatase (pfam PF01725, KomB), and a SIR2-like domain-containing protein (pfam PF13289, KomC) (Fig. 1B, fig. S1 and fig. S2). Notably, SIR2 proteins are key components of immune systems across all three domains of life (*15-19*). These observations urged us to hypothesize that Kongming represents a previously uncharacterized defense system. To investigate the possible anti-phage activity, we cloned the three-gene operon from *Escherichia coli* NCTC13216 under the control of its native promoter into the pET28a plasmid. Expression of Kongming in *E. coli* MG1655 challenged with a panel of coliphages revealed strong immunity against a remarkably wide range of phages (Fig. 1C and fig. S3). Next, we sought to investigate whether Kongming provides immunity at the population level, a defense strategy that only protects bacterial culture at a low multiplicity of infection (MOI), i.e. the ratio of phages to host cells less than 1 (*20*). We infected the liquid cultures expressing Kongming with phage Rao1 at MOI of 3 and 0.3, respectively. The results show that Kongming rescued culture growth only at a MOI of 0.3, whereas infection at an MOI of 3 resulted in a premature growth retardation (Fig. 1D and fig. S4). The data suggest that Kongming provides population-level immunity via triggering cell death or metabolic arrest.

**Fig. 1.**
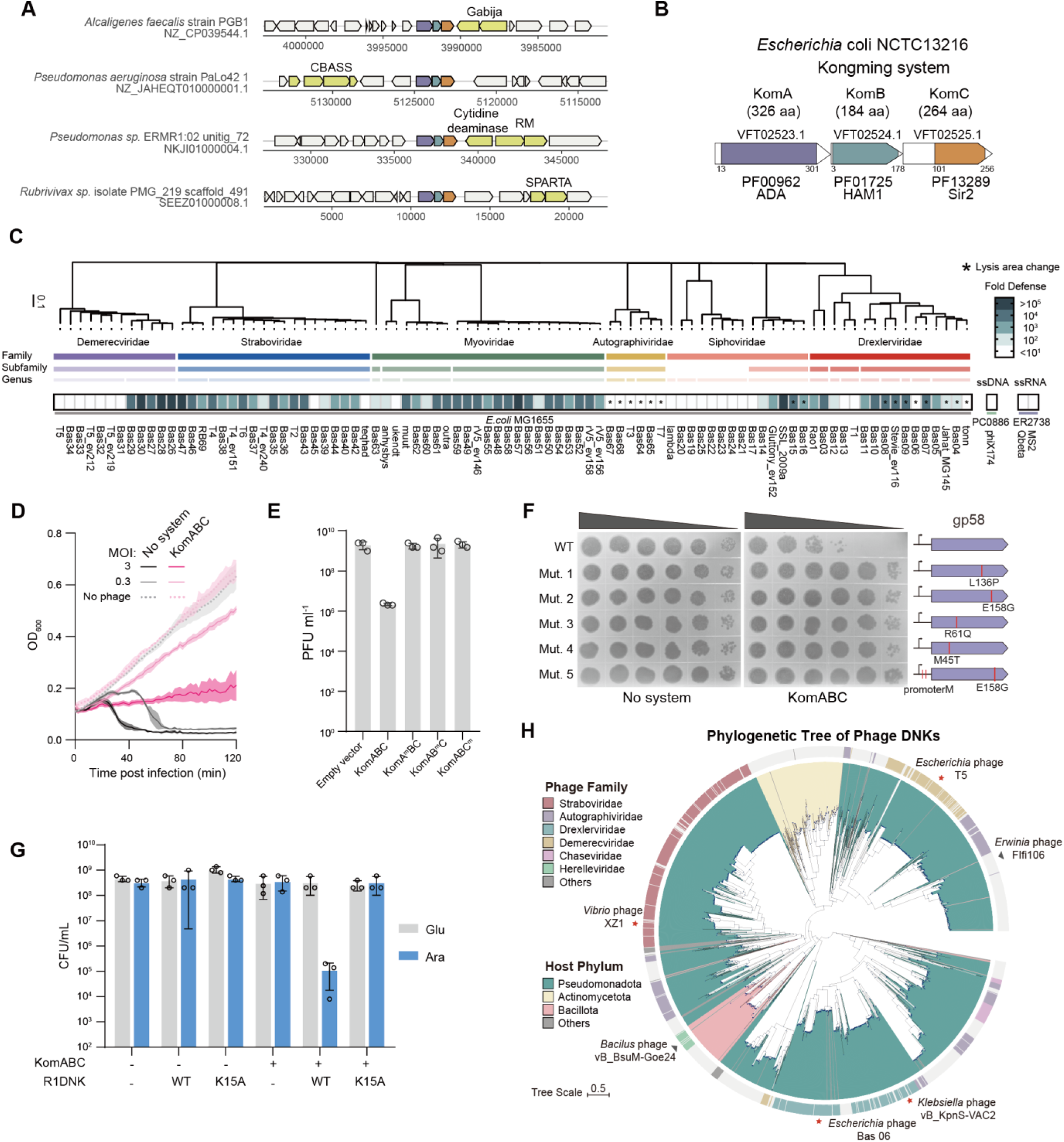
Kongming is an anti-phage system that senses phage DNK. (A) A conserved three-gene operon is found in the neighborhood of known defense systems. Representative examples of the three-gene operon and their genomic contexts are shown. Known defense systems are highlighted in yellow. RM, restriction modification; CBASS, Cyclic oligonucleotide-based antiphage signaling system; SPARTA, short prokaryotic Argonaute TIR-APAZ system. The encoding strain and the respective DNA scaffold accession on the GenBank database are indicated. (B) Domain organization of the three-gene operon from *Escherichia coli* NCTC13216. ADA, adenosine deaminase domain; HAM1, HAM1-like non-canonical purine NTP pyrophosphatase domain. (C) Anti-phage activity of Kongming against a panel of 93 coliphages, showing the mean of three replicates. The branch lengths of the phage phylogenetic tree are scaled using the GBDP formula D0. Horizontal bars represent phages of the same taxonomic group up to the family level. (D) Growth curves of Kongming-expressing *E. coli* MG1655 (KomABC) and control strain (no system) post infection with the Rao1 phage at MOIs of 3 and 0.3. Data represent the mean ±standard deviation of n = 3 biological replicates. (E) Plaque forming units (PFUs) of Rao1 infecting *E. coli* MG1655 strains expressing wild type (WT) or mutated KomABC (A^m^, KomA^H15A-H17A^; B^m^, KomB^N9A-K12A-E15A^; C^m^, KomC^H146A^). Data represent the mean ±standard deviation of n = 3 biological replicates, with individual data points overlaid. (F) Rao1 mutants capable of escaping Kongming immunity carry mutations within the *gp58* gene. PFUs were analyzed for Rao1 and its escaper mutants on *E. coli* MG1655 expressing an empty vector (no system) or KomABC. Representative images of three independent replicates are shown. The mutations are represented on the right. Detailed information about the mutations is listed in table S5. (G) R1DNK triggers cell death in the presence of KomABC. Colony forming units (CFUs) were measured for cells co-expressing KomABC and R1DNK or R1DNK^K15A^ mutant on the plates containing glucose (Glu, to repress R1DNK expression) or arabinose (Ara, to induce R1DNK expression). Data represent the mean ±standard deviation of n = 3 biological replicates, with individual data points overlaid. (H) Phylogenetic analysis of phage DNKs retrieved from the RefSeq complete phage genomes database (INPHARED). The names of phages encoding the experimentally tested DNKs in this work are indicated. Red star, co-expression of DNK and KomABC is toxic; black triangle, not toxic.

To investigate whether each of the three components is essential for anti-phage immunity, we assayed mutated versions of the system. Alanine substitution of the conserved residues within the active site pocket of KomA (His^15^, His^17^) (*21*) and KomC (His^146^) (*22*) (fig. S2) abolished the immune defensive phenotype against Rao1 (Fig. 1E, fig. S5), indicating that the enzymatic activities of KomA and KomC are essential for Kongming defense. In contrast, the conserved catalytic aspartic acid, responsible for nucleophilic attack in experimentally characterized HAM1 family enzymes (*23*), is replaced by histidine in KomB (His^65^) (fig. S2), suggesting that KomB may be catalytically inactive. Nevertheless, KomB retains the conserved residues that are implicated in binding to non-canonical purine NTP substrates in its experimentally characterized homologs (*23*) (fig. S2). Alanine substitution of three substrate-binding residues (Asn^9^, Lys^12^, and Glu^15^) also abrogated immunity (Fig. 1E, fig. S5), underscoring that the binding of KomB to non-canonical purine NTP is required. Together, the data indicate that Kongming is a novel anti-phage system, where each of the three components plays an indispensable role.

### Phage DNKs trigger Kongming immunity

Phages often evolve resistance to bacterial immune defenses by acquiring mutations in the components sensed by the immune systems, thereby providing valuable insights into the system’s activation mechanism (*24, 25*). To identify potential phage-encoded triggers, we infected *E. coli* cells expressing Kongming with Rao1 and isolated immune escaper mutants for further examination (Fig. 1F). Comparative genomics analysis of escaper mutants and the parental Rao1 phage revealed that the escapers all carried mutations within the coding sequence or promoter region of the gene *gp58* (Fig. 1F, table S5), which encodes a phage deoxynucleotide monophosphate kinase (hereafter, R1DNK)(fig. S6A). To investigate whether R1DNK acts as the trigger of Kongming, we analyzed the cell viability and culture growth kinetics of cells co-expressing R1DNK and Kongming (Fig. 1G and fig. S7A). The results show that the co-expression induced extensive cell death and growth suppression, suggesting that R1DNK alone triggers Kongming toxicity. Furthermore, the toxicity was eliminated when the conserved catalytic residue Lys^15^ of R1DNK was mutated to alanine (K15A), indicating that the enzymatic activity of R1DNK is essential for triggering the Kongming immunity (Fig. 1G, fig. S7A-B and fig. S5).

Phage DNKs can catalyze the phosphorylation of dNMP or modified dNMP to the corresponding dNDP, which are then converted to dNTPs for rapid phage genome replication (fig. S6A) (*26, 27*). To investigate the function of R1DNK, we constructed a *dnk* deletion Rao1 mutant (Rao1^Δdnk^) using CRISPR-Cas13 (*28*). The mutant was resistant to Kongming, reinforcing that R1DNK is the trigger (fig. S8A). Infection with Rao1 induced a gradual ∼2-3 folds increase in the dAMP, dADP and dATP levels (fig. S8B). Deletion of *dnk* resulted in a moderately lowered dATP level, indicative of a role of R1DNK in phage dNTP systhesis (fig. S8B). Nevertheless, the phage competition assay indicates that deletion of *dnk* did not cause apparent fitness cost, implying that the DNK-mediated dNTP synthesis is not critical for Rao1 replication under our experimental conditions (fig. S8C).

Phylogenetic analysis revealed that DNKs are widely spread across phages pertaining to distinct families (Fig. 1H, fig. S7C), consistent with the remarkably broad protective activity of Kongming. To test if Kongming is activated by other phage DNKs, we selected six representative DNK homologs from different branches of the phylogenetic tree and co-expressed them with Kongming. These results revealed that the DNKs from T5 phage, Klebsiella phage vB_KpnS-VAC2, Vibrio phage XZ1, and Escherichia phage KarlJaspers triggered Kongming toxicity (Fig. 1H and fig. S7D). In contrast, the DNK form Bacillus phage vB_BsuM-Goe24 or Erwinia phage FIfi106 did not induce Kongming-mediated cell death (Fig. 1H and fig. S7D), suggesting that these DNKs either have distinct enzymatic properties or we could not recapitulate their activity through heterologous expression in *E. coli*. Interestingly, we found that the distribution of DNKs in our tested phage collection correlates well with the Kongming immunity, except for a minority of phages (fig. S3). These results support the notion that Kongming is triggered by phage-encoded DNKs or their enzymatic activities. Notably, within the exceptions of DNK-encoding phages, T5-like phages are particularly resistant to Kongming (Fig. 1C). However, expression of T5DNK in the absence of infection robustly activates Kongming toxicity (fig. S7D), strongly suggesting that T5-like phages encode anti-Kongming factors. On the other hand, a few phages that are sensitive to Kongming do not encode DNK (fig. S3), indicating that Kongming might also be triggered through alternative mechanisms that require further investigation.

### KomB and KomC form an effector complex capable of NAD^+^ degradation

In various SIR2-containing anti-phage systems, such as Thoeris, defence-associated sirtuins (DSRs) and SIR2-containing pAgo systems, SIR2 acts as the cell-killing effector module via rapid cellular NAD^+^ depletion (*12, 29, 30*). To test whether KomC similarly degrades NAD^+^, we measured NAD^+^ levels in the cells co-expressing R1DNK and Kongming (Fig. 2A). Indeed, co-expression resulted in significant NAD^+^ depletion, and mutation in the active site of KomC abrogated this activity, confirming that KomC is responsible for NAD^+^ degradation. The NAD^+^ depletion was accompanied by extensive cell death (fig. S7E), in line with our previous observations (Fig. 1G). Further, the observed NAD^+^ depletion was also abrogated when KomA, KomB, or R1DNK were mutated in their respective conserved sites, indicating that the activation of KomC is dependent on the activities of KomA and R1DNK and the substrate-binding ability of KomB.

**Fig. 2.**
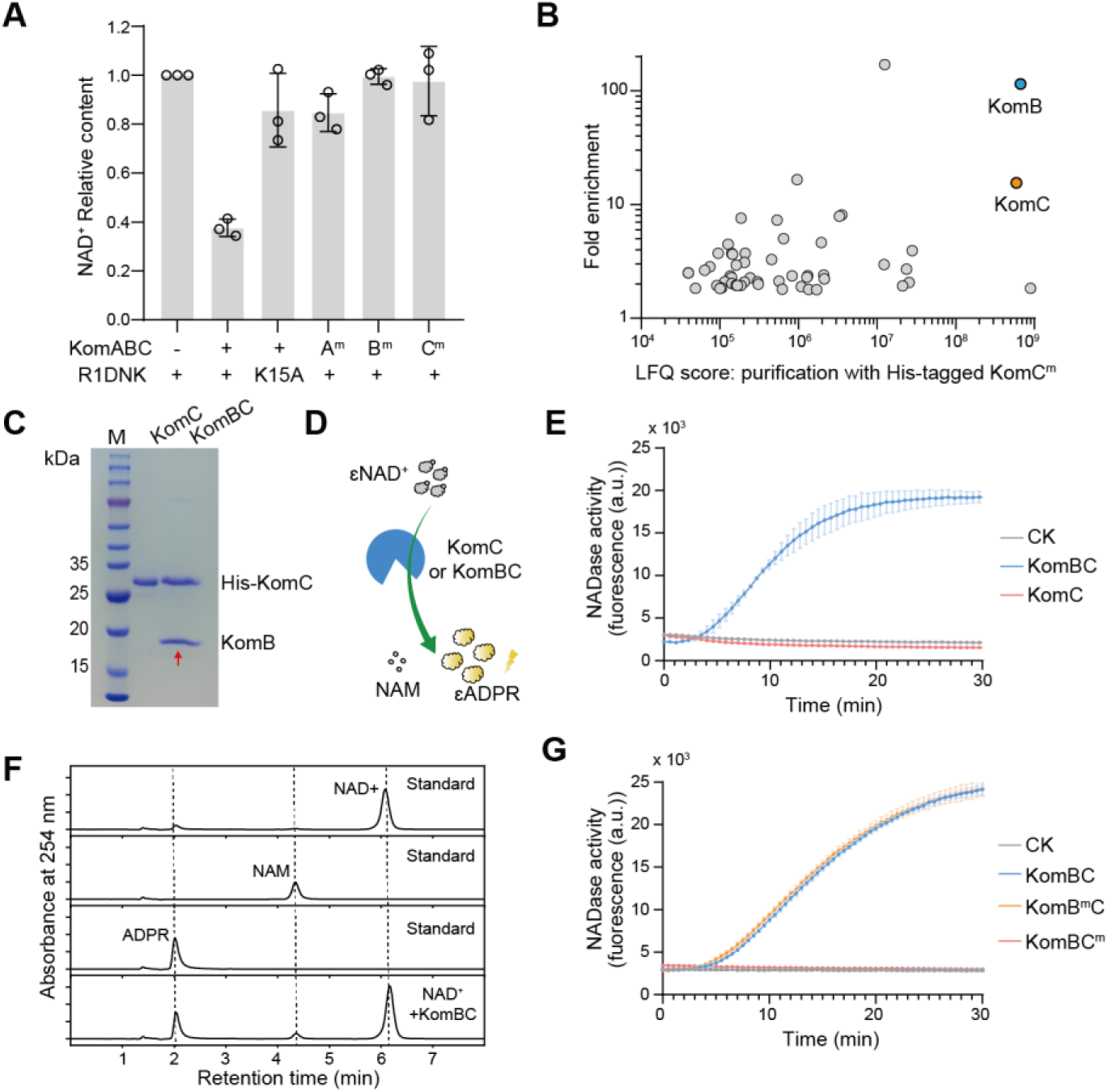
KomB and KomC form an effector complex that mediates NAD^+^ depletion. (A) R1DNK triggers cellular NAD^+^ depletion in the presence of KomABC. NAD^+^ levels were measured for cells co-expressing KomABC and R1DNK, or their mutants: A^m^, KomA^H15A-H17A^; B^m^, KomB^N9A-K12A-E15A^; C^m^, KomC^H146A^. Data represent the mean ±standard deviation of n = 3 biological replicates, with individual data points overlaid. (B) Mass spectrometry of the proteins purified from cells expressing KomA, KomB, His-tagged KomC^H146A^ and R1DNK. Shown are the LFQ scores (X-axis) and enrichment folds (Y-axis) compared to the sample purified from cells expressing KomA, KomB, untagged KomC^H146A^ and R1DNK. Only the significantly enriched proteins are shown. KomB and KomC are indicated in the panel. (C) SDS-PAGE analysis of purified His-KomC and the KomB-(His)KomC complex (KomBC). The protein band indicated by the red arrow was validated to be KomB by mass spectrometry. (D) Schematic of the fluorogenic reaction used to analyze the NADase activity of KomBC and KomC. (E) NADase activity of purified 0.8 μM KomC and KomBC with ε-NAD as substrate. CK, reaction without any enzyme. Data represent the mean ±standard deviation of n = 3 biological replicates. (F) High-performance liquid chromatography (HPLC) analysis of the NAD^+^ degradation products of 0.1 μM KomBC, and NAD^+^, ADPR, and NAM as standards. (G) NADase activity of 0.6 μM KomBC and its mutants: B^m^, KomB^N9A-K12A-E15A^; C^m^, KomC^H146A^. CK, reaction without KomBC. Data represent the mean ±standard deviation of n = 3 biological replicates.

To investigate whether KomC activation requires effector complex formation, we examined its potential interactions with KomA, KomB, or R1DNK. To this end, we co-expressed R1DNK and Kongming, where KomC carried a 6XHis tag to allow Ni-NTA affinity purification and a H146A mutation to abolish its toxic NADase activity (fig. S9A). The co-purified proteins were subjected to mass spectrometry (MS) analysis to query the possible interaction with KomC. Compared to the control purification with untagged KomC^H146A^, KomB and KomC were significantly enriched in the purification with His-tagged KomC^H146A^ (Fig. 2B and fig. S9B, table S6), indicative of a direct physical interaction between KomB and KomC. SDS-PAGE and MS analysis confirmed that both KomB and KomC were retrieved by Ni-NTA affinity purification from cells co-expressing KomB and His-tagged KomC, reinforcing the notion that the two proteins form a stable complex (Fig. 2C and fig. S9C, table S7). Additionally, pull-down experiments using MBP-tagged KomA and Strep-tagged R1DNK did not detect any interaction between KomA and KomBC complex or between the phage-encoded trigger and the three proteins of Kongming (fig. S9D and E), suggesting that R1DNK activates Kongming indirectly rather than through protein-protein interaction.

We then analyzed whether KomC, standalone or in the complex with KomB, has NADase activity using ε-NAD as substrate, the degradation of which produces a fluorescent product (Fig. 2D). The KomBC complex efficiently degraded ε-NAD in the presence of Zn^2+^, whereas KomC alone did not cleave ε- NAD in the same condition (Fig. 2E, fig. S10A), underscoring the requirement of complex formation with KomB for the NADase activity. Analysis of the NAD^+^ degradation products indicates that KomBC degraded NAD^+^ into NAM and ADPR, in line with the products of the SIR2 effectors in other defense systems (*12, 29*) (Fig. 2F). Further, the NADase activity was abrogated by the mutation of the KomC active site but not affected by the mutation of KomB (fig. S10B and Fig. 2G), confirming that the active site of KomC catalyzes the degradation of NAD^+^. Taken together, these results demonstrate that KomB and KomC form the effector complex that is responsible for immune NAD^+^ depletion.

### dITP activates the KomBC effector complex

The observation that neither R1DNK nor KomA physically interact with the KomBC complex, despite their activities being essential for triggering NAD^+^ degradation, led us to speculate that KomA and R1DNK activate KomBC via synthesizing a second messenger. The predicted functions of KomA, R1DNK, and KomB are linked to nucleotide metabolism pathways, suggesting that Kongming could rely on a previously unrecognized nucleotide-based immune messenger. Given that KomB forms a stable complex with KomC (Fig. 2C) and has a mutated catalytic site (fig. S2), we hypothesized that KomB may function as a signal recognition module. Additionally, previously characterized KomB family proteins, known as inosine triphosphate pyrophosphatases (ITPases), are house-keeping enzymes that hydrolyze noncanonical nucleoside triphosphates to prevent their incorporation into nascent DNA and RNA chains, e.g. degrading ITP and dITP to IMP and dIMP, respectively (*23, 31*). Together, these observations suggest that the putative signaling molecule could be a noncanonical nucleotide.

To test this idea, we analyzed the NADase activity of KomBC in the presence of dITP, ITP, IDP, IMP, dUTP, or canonical nucleotides (dNTPs and NTPs). The results show that dITP activates the NADase activity of KomBC most robustly (Fig. 3A). While IDP also activated KomBC, it triggered a >100-fold weaker response compared to dITP, and the other tested nucleotides did not activate KomBC (Fig. 3A and fig. S10C-D). Upon activation, the NADase activity generated the same products as those in the absence of dITP (fig. S10E). These findings indicate that dITP is the signaling molecule that activates KomBC for NAD^+^ degradation. Furthermore, the KomBC complex with inactive KomC did not exhibit any NADase activity, whereas the KomB mutation still retained the basal activity but abolished the dITP-stimulated activity (Fig. 2G and Fig. 3B). To test whether KomBC degrades the dITP immune signal, which may represent an autoinhibition mechanism observed in type III CRISPR-Cas system signaling pathways (*32*), we incubated dITP with KomBC and analyzed the potential products (fig. S10F). However, no products were detected, indicating that KomB has lost its dITP pyrophosphatase activity. Together, these results indicate that KomB functions as the signal-recognition module that senses dITP as an immune signal and mediates activation of KomC.

**Fig. 3.**
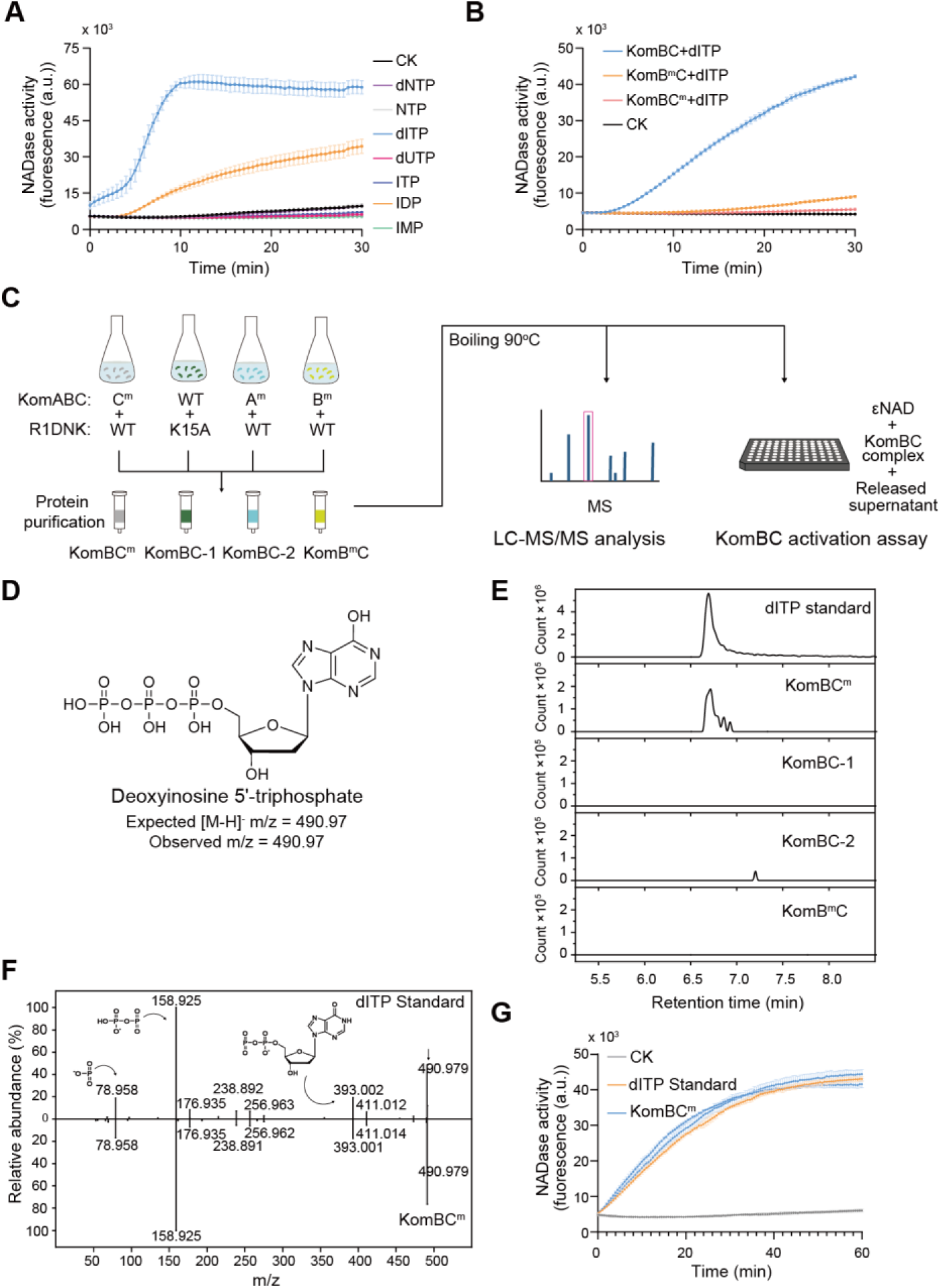
dITP signaling activates the KomBC effector complex. (A) NADase activity of 50 nM KomBC in the presence of the indicated nucleotides. CK, reaction without any nucleotide. Data represent the mean ±standard deviation of n = 3 biological replicates. (B) NADase activity of 50 nM KomBC and its mutants in the presence of dITP. CK, reaction without KomBC. Data represent the mean ±standard deviation of n = 3 biological replicates. (C) Schematic representation of the experimental setup. The cells expressing mutated KomABC (A^m^, KomA^H15A-H17A^; B^m^, KomB^N9A-K12A-E15A^; C^m^, KomC^H146A^) and wild type R1DNK, or wild type KomABC and mutated R1DNK, were used for purification of different KomBC complexes. The complexes were subsequently denatured, and the released supernatants were analyzed by LC-MS/MS and tested for their ability to activate the NADase activity of wild type KomBC. (D) Chemical structure of dITP and its m/z value. (E) LC-MS analysis of the released supernatants from the denatured protein samples from (C). Extracted mass chromatograms of ions with an m/z value of 490.97 are presented. Shown are the representative data of two independent replicates. (F) MS/MS analysis of the released molecules from KomBC^m^, using dITP as a standard. (G) NADase activity of 50 nM KomBC in the presence of the supernatant from KomBC^m^, where the concentration of dITP was diluted to 1 nM. The experiments were performed in two independent replicates. dITP standard: 50 nM KomBC with 1 nM synthetic dITP; CK: 50 nM KomBC.

### Kongming and R1DNK produce dITP signaling molecules *in vivo*

Next, we aimed to analyze whether Kongming synthesizes the signaling molecule dITP in response to phage infection *in vivo*. We hypothesized that the synthesized signal could be bound by KomBC. Therefore, to retrieve the putatively bound molecule, we purified the KomBC^m^ complex from cells expressing His-tagged KomC^m^, KomA, KomC, and the phage trigger R1DNK (Fig. 3C and fig. S11A). The purified KomBC^m^ complex was subsequently denatured and precipitated, and the resulting supernatant was analyzed by liquid chromatography-tandem mass spectrometry (LC-MS/MS) to investigate whether it contains dITP (Fig. 3D-F). The analysis detected a molecule that has the same retention time, m/z value, and molecular fragments as dITP. In addition, the molecule was absent in the KomBC complex purified from the cells containing a mutated KomA or R1DNK, or in the KomB^m^C complex from the cells containing wild-type KomA and R1DNK (Fig. 3E). The data suggest that KomBC binds to dITP as a signaling molecule and that the enzymatic activities of both KomA and R1DNK are required for dITP synthesis *in vivo*.

To explore whether the dITP in the supernatant activates KomBC, we measured the concentration of the released dITP via LC-MS/MS (fig. S11B), diluted it to the same dITP concentation as the standard, and analyzed its capability to activate KomBC (Fig. 3G). The results revealed similar activation efficiency of the diluted supernatant and the dITP standard. In addition, the supernatant from the cultures with mutation in KomA, R1DNK or KomB failed to activate KomBC (fig. S11C). Together, these results confirm that dITP is the sole signaling molecule synthesized by Kongming in response to phage DNK and recognized by KomBC for immune activation.

### Biosynthetic pathway of the dITP immune signal

To reveal how dITP is synthesized during the immune response, we started to characterize the biochemical properties of KomA and R1DNK, which are shown essential for dITP synthesis (Fig. 3E). Given that characterized homologs of KomA catalyze deamination of adenosine and deoxyadenosine to inosine and deoxyinosine, respectively (Fig. 4A) (*33*), we tested the substrate specificity of KomA against diverse adenine nucleotides. Incubation with dAMP, dADP, dATP, AMP, ADP, or ATP, followed by the analysis of the resulting products via HPLC revealed that KomA shows the highest adenosine deaminase activity with dAMP as a substrate and produces dIMP, whereas the deamination of dADP, AMP, and ADP were also observed (Fig. 4B and fig. S12). Nevertheless, deamination of dATP was not detected (Fig. 4B), suggesting that KomA cannot complete dITP production by itself, which possibly requires a factor that becomes available during phage infection.

**Fig. 4.**
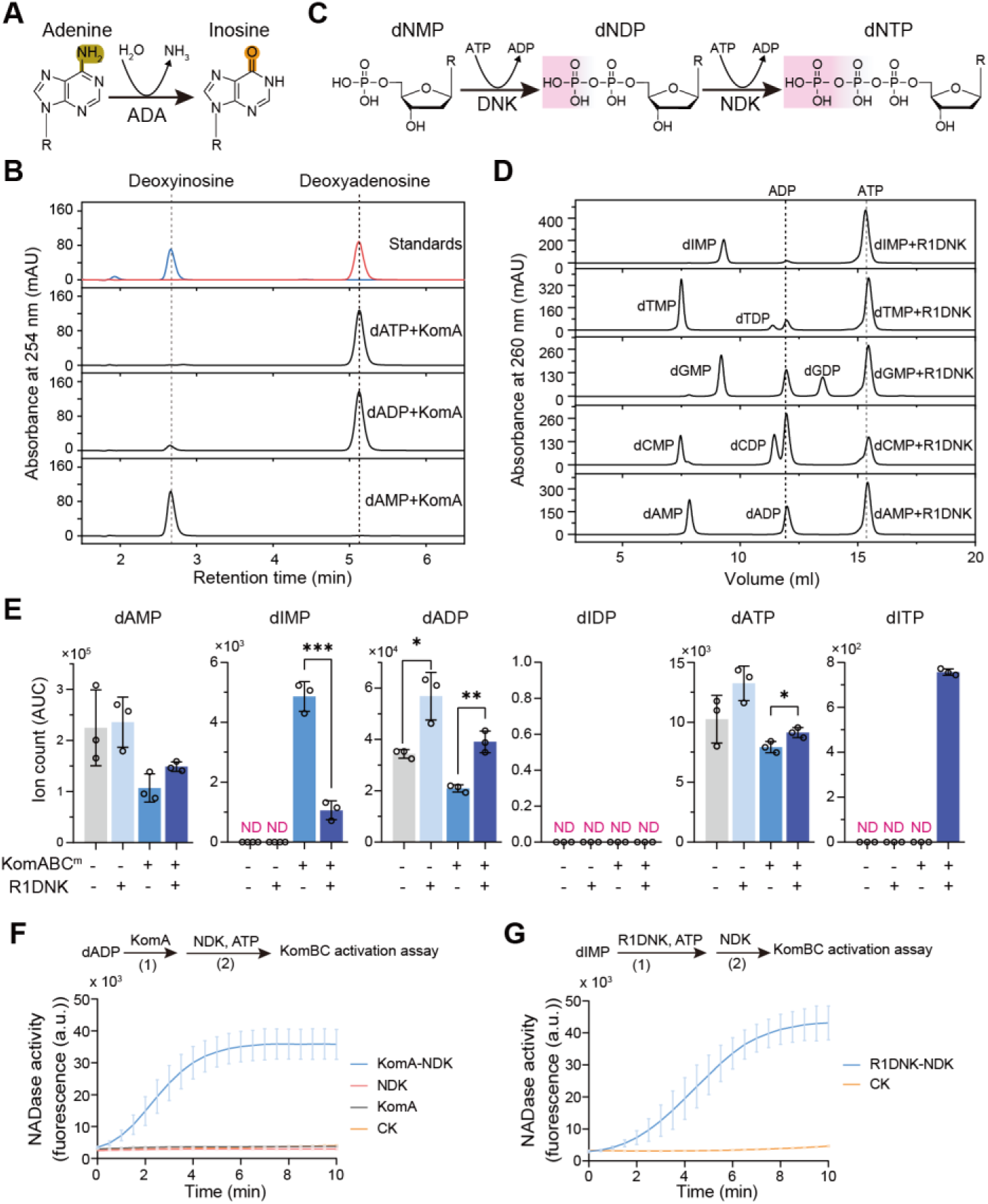
Synthesis of the anti-phage signal dITP. (A) Schematic of the deamination reaction by typical adenosine deaminase enzymes (ADA). (B) High-performance liquid chromatography (HPLC) analysis of the reactions of KomA using dATP, dADP or dAMP as substrates. Deoxyadenosine and deoxyinosine were also analyzed by HPLC as standards. (C) Schematic of the phage dNTP synthesis pathway performed by phage DNK and host NDK, using dNMP as precursors. (D) Ion-exchange chromatography (IEC) analysis of the reactions of R1DNK with ATP as a phosphate donor and dIMP, dTMP, dGMP, dCMP, or dAMP as substrates. (E) Ion count (area under curve) of dAMP, dIMP, dADP, dIDP, dATP, and dITP in lysates extracted from cells expressing KomABC^m^ and/or R1DNK. Data represent the mean ±standard deviation of n = 3 biological replicates, with individual data points overlaid. *, p<0.05; **, p<0.01; ***, p<0.001. ND, not detected. (F) NADase activity of 50 nM KomBC in the presence of the reaction product of KomA and NDK with dADP as substrate and ATP as phosphate donor. The reaction mixtures containing only KomA or NDK were also tested. CK: NADase activity of KomBC in the presence of dADP and ATP. Data represent the mean ±standard deviation of n = 3 biological replicates. (G) NADase activity of 50 nM KomBC in the presence of the reaction product of R1DNK and NDK with dIMP as substrate and ATP as phosphate donor. CK: NADase activity of KomBC in the presence of dIMP and ATP. Data represent the mean ±standard deviation of n = 3 biological replicates.

Therefore, we hypothesized that Kongming hijacks the phage DNKs to produce dITP as a triggering mechanism. Phage DNKs are supposed to phosphorylate dNMP to dNDP (fig. S6A)(*26, 27*), which are then converted to dNTPs by host nucleoside diphosphate kinases (NDKs)(*34*) to faciliate phage genome replication (Fig. 4C). Biochemical characterization of R1DNK show that R1DNK efficiently phosphorylated dAMP, dGMP and dCMP to dADP, dGDP, and dCDP, respectively, and also showed kinase activity with dTMP as a substrate (Fig. 4D and fig. S13). However, the conversion of dIMP to dIDP was minimal (Fig. 4D). Together, the data suggest that KomA and R1DNK can catalyze a cascade reaction to produce dIDP from dAMP.

To explore how KomA and R1DNK produce dITP *in vivo*, we analyzed the levels of adenine nucleotides and their inosine derivatives in cells expressing R1DNK and Kongming carrying a KomC^H146A^ mutation (KomABC^m^). The cells expressing KomABC^m^ accumulated dIMP but not dIDP or dITP, indicating that of KomA only deaminates the host dAMP (Fig. 4E). Expression of R1DNK resulted in elevated dADP level, confirming its capability to phosphorylate dAMP (Fig. 4E). In the cells co-expressing R1DNK and KomABC^m^, dITP, the final signaling molecule, was accumulated, while dIDP was still not detectable (Fig. 4E). These results suggest that dIDP can be rapidly converted to dITP. In addition, compared to expression of KomABC^m^ alone, co-expression of R1DNK and KomABC^m^ resulted in lowered dIMP level, implying that R1DNK may phosphorylate dIMP *in vivo*.

To investigate whether host NDK can convert dIDP to dITP, we sought to reconstitute the cascade reaction of dITP production. We firstly incubated dADP with KomA, and then with *E. coli* NDK and ATP, and analyzed the product for its ability to activate KomBC (Fig. 4F). As expected, the product efficiently activated KomBC for NAD^+^ degradation (Fig. 4F). LC-MS/MS analysis confirmed that the product contained dITP (fig. S14A-B). In addition, the absence of either ADA or NDK abolished dITP generation, indicating that NDK is a host enzyme involved in dITP synthesis (Fig. 4F and fig. S14A). Next, we incubated dIMP with R1DNK and ATP, and then with NDK. The resulting product also activated KomBC (Fig. 4G). Further, using LC-MS/MS, we detected dIDP as the product after incubation of dIMP with R1DNK and ATP (fig. S14C-E). The data indicate that R1DNK indeed harbors the dIMP phophorylation activity, although LC failed to detect the product (Fig. 4D). Collectively, the results establish that KomA and R1DNK can generate dIDP from dAMP and that host NDK converts the generated dIDP to dITP as the signaling molecule. The dIDP can be generated via two pathways (fig. S14F): (i) phosphorylation of dAMP by R1DNK followed by deamination of dADP by KomA, and (ii) dAMP deamination by KomA followed by phosphorylation of dIMP by R1DNK. Both the two pathways may contribute to immune signaling *in vivo*, while the specific extent of the contribution of each pathway might be dependent on the phage DNK substrate specificity and nucleotide metabolism mechanism.

### Phage-encoded inhibitor of Kongming immunity

Our observation that T5 is resistant to Kongming, despite the robust triggering of Kongming by T5DNK expression, led us to hypothesize that T5 encodes an anti-Kongming factor (Fig. 1C and fig. S7D). In other anti-phage signaling systems, such as CBASS, Thoeris, and type III CRISPR-Cas, phages typically counteract immunity by targeting the signaling molecule, either by sponging it or leading to its degradation (*35-38*). We therefore hypothesized that T5 may encode an anti-Kongming factor with dITP-binding or -processing ability. Previous studies have shown that T5 encodes a deoxyribonucleotide 5’ monophosphatase (T5Dmp) that converts 5’-dNMP to deoxynucleosides before phage genome replication begins (*39*), implying that it may subvert Kongming immunity by degrading dAMP, the precursor for dITP synthesis. To confirm this, we constructed a *dmp* deletion T5 mutant (T5^Δdmp^) and analyzed the sensitivity of T5^Δdmp^ to Kongming. Compared to wild-type T5, T5^Δdmp^ could not overcome Kongming immunity, indicating that T5Dmp indeed functions as an anti-Kongming factor (Fig. 5A). Analysis of nucleotides levels post infection shows that wild type T5 resulted in lowered dAMP level at 5 min, while T5^Δdmp^ induced a sharp accumulation of dAMP (Fig. 5B), confirming that T5 infection degrades the host genome into dNMP, while T5Dmp is responsible for depletion of cellular dNMP (*39*). Moreover, dAMP accumulation post T5^Δdmp^ infection was accompanied with elevated dADP and dATP levels, indicative of efficient phosohorylation of dAMP to these nucleotides (Fig. 5B).

**Fig. 5.**
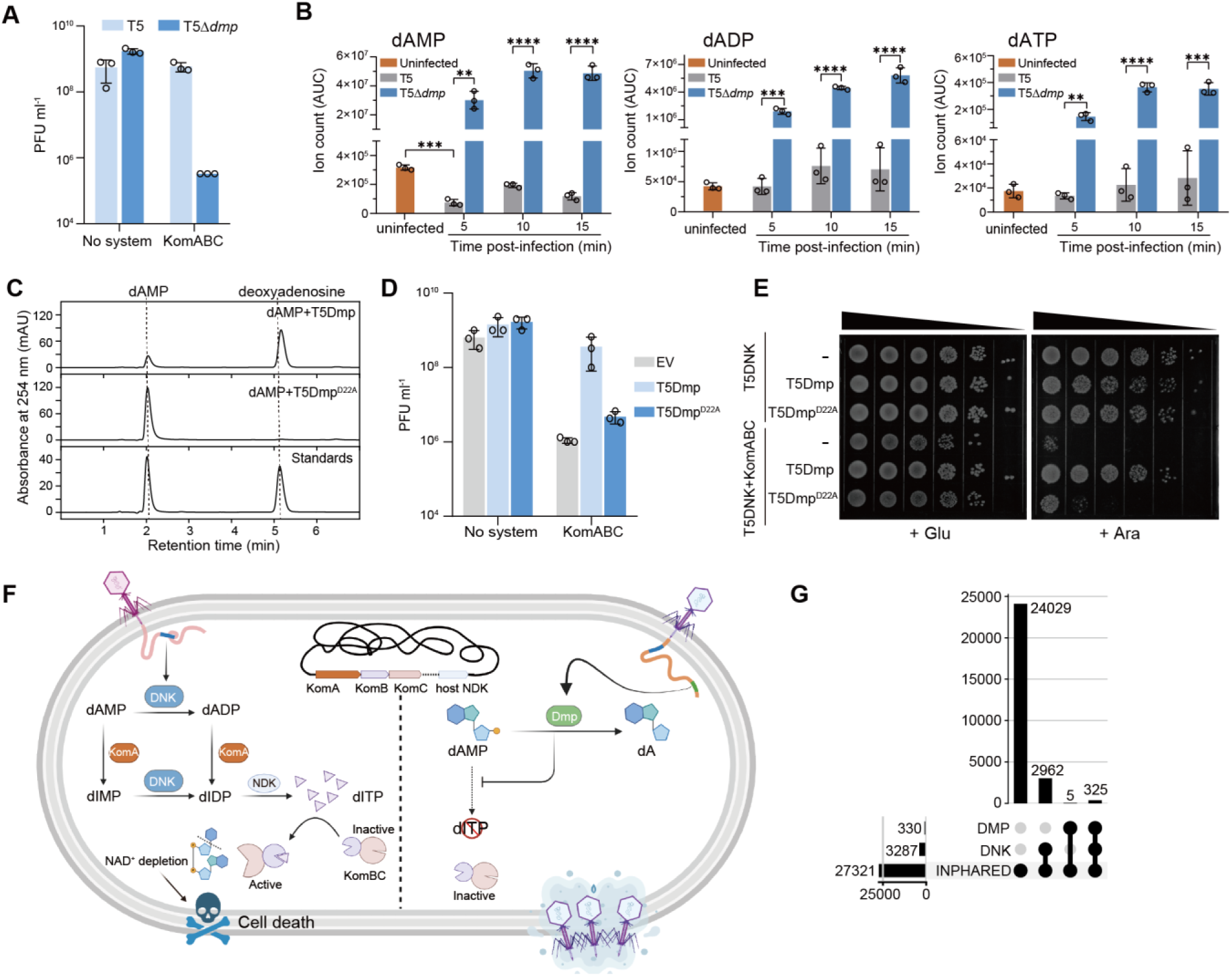
T5 5’-deoxynucleotidase inhibits Kongming by degrading dAMP. (A) Deletion of *T5dmp* renders T5 sensitive to Kongming. Plaque forming units (PFU) were analyzed for T5 and the *dmp* deletion mutant on *E. coli* MG1655 lawns with or without KomABC. Data represent the mean ±standard deviation of n = 3 biological replicates, with individual data points overlaid. (B) Ion count (area under curve) of dAMP, dADP, and dATP in lysates extracted from cells infected with T5 and T5^Δdmp^, or uninfected cells. Data represent the mean ±standard deviation of n = 3 biological replicates, with individual data points overlaid. **, p<0.01; ***, p<0.001; ****, p<0.0001. (C) Mutation at the putative active site abolishes the dAMP phosphatase activity of T5Dmp. dAMP was incubated with T5Dmp or a D22A mutant, and the product was analyzed with HPLC. Deoxyadenosine and dAMP standards are shown at the bottom. (D) The D22A mutation abolishes the anti-Kongming function of T5Dmp. Cells expressing KomABC or not were transformed with an empty vector (EV) or plasmids encoding T5Dmp or its D22A mutant. Then, PFUs were analyzed for Rao1 infecting the bacterial lawns of the transformants. Data represent the mean ±standard deviation of n = 3 biological replicates, with individual data points overlaid. (E) T5Dmp inhibits the T5DNK-triggered toxicity in the cells expressing KomABC. Bacterial viability was measured for cells expressing KomABC, T5DNK, and T5Dmp or its D22A mutant on agar plates containing glucose (Glu, to repress T5DNK expression) or arabinose (Ara, to induce T5DNK expression). (F) Left: the proposed model for the Kongming immune pathway. Phage DNK and KomA, together with host NDK, produce dITP as the signaling molecule using dAMP as precursor; then dITP activates the KomBC complex for NAD^+^ depletion, resulting in death of the infected cells. Right: the anti-Kongming function of phage-encoded Dmps. Phage Dmp degrades dAMP to dA (deoxyadenosine), thus inhibiting dITP synthesis. Created with BioRender.com. (G) Correlation of phage-encoded DNKs and Dmps in phage genomes retrieved from the INPHARED database.

We further examined the phosphatase activity of T5Dmp and, in line with previous studies (*40*), T5Dmp displayed phosphatase activity with dNMP substrates, as well as dIMP and dADP, but failed to degrade dITP (fig. S15A-B). These results indicate that the anti-Kongming function of T5dmp is not achieved by the degradation of the signaling molecule. To confirm the dependency of T5Dmp’s anti-Kongming function on its enzymatic activity, we constructed a T5Dmp mutant that carries an alanine substitution at D22, a conserved residue in phosphatases that functions in Mg^2+^ coordination (fig. S15C) (*41*). Indeed, compared to wild-type T5Dmp, the D22A mutant showed no phosphatase activity with dAMP as substrate (Fig. 5C). Consistently, the D22A mutation abolished the ability of T5Dmp to protect Rao1 from Kongming immunity, suggesting that T5Dmp’s phosphatase activity is required for its anti-defense function (Fig. 5D). To investigate whether T5Dmp inhibits Kongming by interfering the DNK-triggered signaling pathway, we cloned T5Dmp into the cells co-expressing T5DNK and Kongming. Here, T5Dmp suppressed the Kongming immunity-triggered cell death, while the D22A mutation restored the immunity (Fig. 5E). Together, our data collectively indicate that T5Dmp is a robust anti-Kongming inhibitor that functions via removal of cellular dAMP, the precursor for dITP synthesis (Fig. 5F).

To further explore the functional impact of T5Dmp and T5DNK, we compared the fitness of wild type T5 and T5 mutants carrying *dmp* and *dnk* deletions via phage competition assays. This revealed that deletion of *dmp* resulted in severe fitsness cost (fig. S16A). Deletion of *dnk* also induced fitness defects in both wild type T5 and T5^Δdmp^ (fig. S16A). The results suggest that T5DNK and T5Dmp play important roles in the phage replication cycle. Notably, T5 encodes its own de novo nucleotide synthesis pathways (*42*), which might involve the DNK activity. In addition, infection with T5^Δdmp-dnk^ still induced accumulation of dADP and dATP, suggesting an alternative dAMP phosphorylatation mechanism (fig. S16B). Consistently, the T5^Δdmp-dnk^ phage was sensitive to Kongming, indicating that DNK is not the sole trigger (fig. S16C). Taken together with the phenotypes of Rao1, our data indicate that T5 and Rao1 encode distinct nucleotide metabolism pathways, with their DNKs likely adapted to different functions (fig. S6).

Phylogenetic analysis of Dmp homologs across phages revealed their frequent co-existence with DNKs (Fig. 5G and fig. S17), reinforcing their function as anti-Kongming proteins that prevent Kongming immune activation by phage DNKs. To further investigate this hypothesis, we selected three additional Dmps and tested their ability to inhibit the toxicity from the co-expression of DNK and Kongming. Two of them originate from distantly-related phages not present in our tested panel (Erwinia phage KEY and Vibrio phage VPG01), and the third one is from Bas26, a phage shown to be sensitive to Kongming immunity (fig. S3). While we were unable to construct the expression plasmid for Erwinia phage KEY Dmp (probably due to its toxicity in *E. coli*), the Dmps from Vibrio phage VPG01 and Bas26 robustly suppressed cell death triggered by Kongming and their respective DNKs (fig. S15D and fig. S17), indicating their anti-Kongming function. The reason why Bas26 is still sensitive to Kongming (fig. S3) given that it encodes a functional anti-Kongming factor is puzzling and requires further investigation.

### Distribution and genetic associations of Kongming components

Given that previously studied HAM1 family proteins are recognized housekeeping enzymes implicated in the removal of non-canonical nucleotides in cells (*23, 31*), we asked whether defense-associated HAM1 proteins represent a specialized subfamily adapted to immune functions. To address this question, we conducted a phylogenetic analysis of a diverse set of 154,151 HAM1 proteins, with representatives from all three domains of life. The analysis revealed that known housekeeping HAM1 proteins are scattered across many clades of the tree, yet excluded from the distinct clade including KomB (hereafter defensive HAM1 clade) (Fig. 6A). Notably, compared to a clade with housekeeping HAM1s (non-defensive clade), members of the defensive HAM1 clade are frequently located near known anti-phage systems and within plasmid and prophage elements, further supporting their immune function (Fig. 6B). Furthermore, the defensive HAM1s invariably carry the histidine substitution of the catalytic aspartic acid residue that is conserved in non-defensive HAM1s (Fig. 6C). On the other hand, the defensive HAM1s retain the conserved non-canonical nucleotide-binding residues that constitute the substrate-binding pocket in the housekeeping HAM1s (Fig. 6C-D and fig. S18). Together, the data suggest that the defensive HAM1 has evolved from active pyrophosphatases to catalytically-dead proteins that function as the signal-sensing module.

**Fig. 6.**
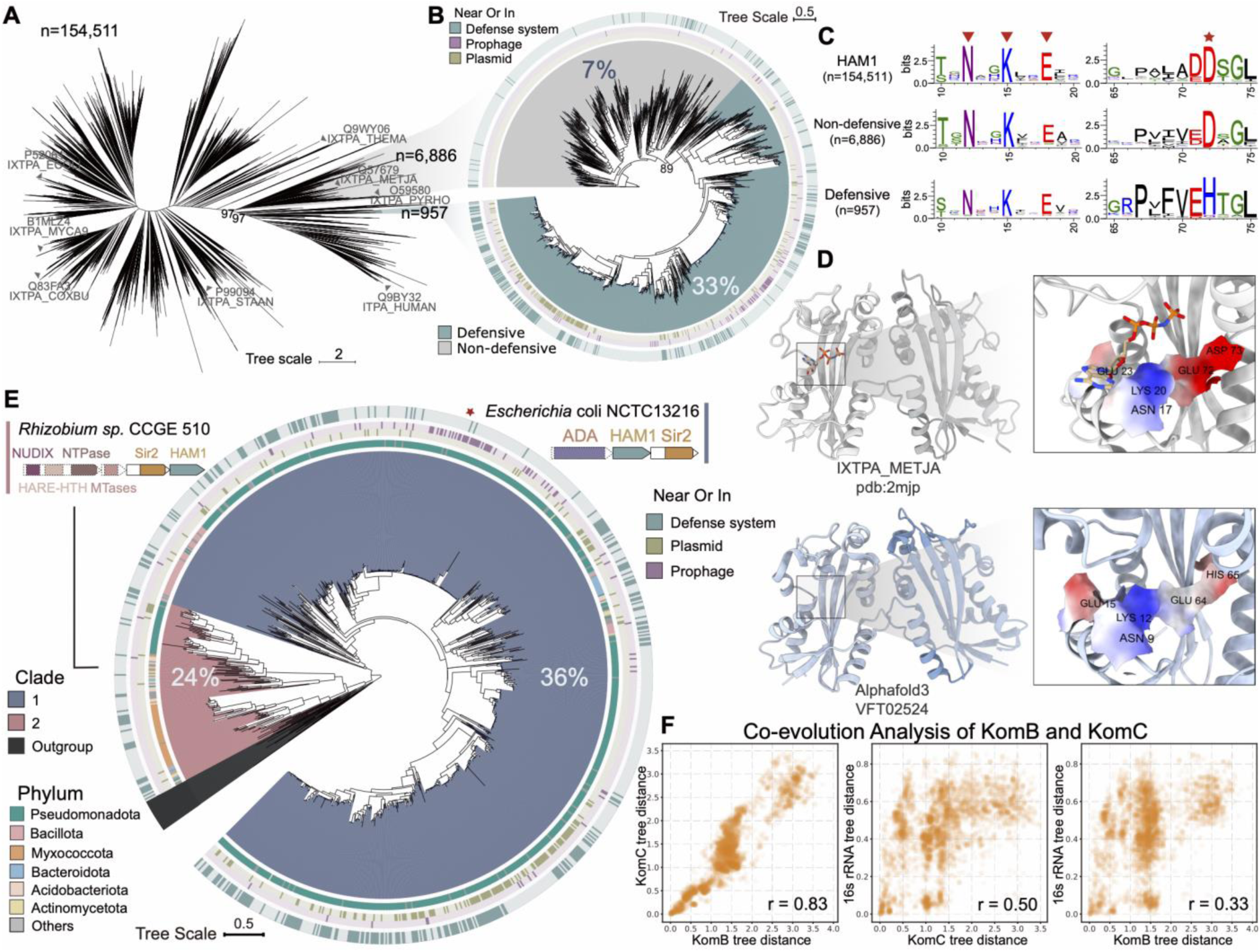
Evolutionary analysis of Kongming components. (A) Phylogenetic tree of HAM1 domain-containing proteins. An unrooted tree was constructed using HAM1 domain-containing proteins from the non-redundant protein database, with sequences longer than 100 amino acids (n=154,511). The sequence numbers of the two highlighted clades (gray and blue) are indicated. Local support values are shown for the indicated branches. Characterized housekeeping HAM1 proteins with experimentally determined structures are indicated with a gray arrow. (B) Phylogenetic tree of defensive and non-defensive HAM1s. All non-redundant sequences of defensive HAM1s were used, while only representative sequences of non-defensive HAM1s were included. For each clade, the percentage of HAM1s encoded near known defense systems is indicated. From outer to inner circles: whether each HAM1 is encoded near known defense systems, near or in prophages, and near or in plasmids. (C) Sequence logo representation of all, non-defensive and defensive HAM1s. Representative regions containing conserved substrate-binding residues (left panel) and the substitution of the catalytic aspartic acid by histidine in defensive HAM1s (right panel) are shown. (D) Structure comparison between a housekeeping HAM1 (IXTPA_METJA, PDB : 2mjp) and the AlphaFold3 structure (ipTM = 0.86, pTM = 0.87) of KomB. A close-up view of the substrate-binding area is shown with electrostatic surfaces of the residues indicated in Fig. 6C. (E) Phylogenetic tree of defensive HAM1s. Twenty sequences of non-defensive HAM1s were used as an outgroup. The percentage of HAM1s encoded near known defense systems is indicated for each clade. Representative operons encoding HAM1 and Sir2 domains are shown. From outer to inner circles: whether each HAM1 is encoded near known defense systems, near or in prophages, near or in plasmids, and the bacterial phylum to which the HAM1 belongs. The red star indicates the HAM1 characterized in this study. (F) Spearman analysis reveals strong co-evolutionary signal of KomB and KomC. The panels show plots of pairwise distances between KomB, the associated KomC, and the 16S rRNA of the organisms encoding the KomB and KomC, respectively. The Spearman rank correlation coefficient is indicated on each plot.

Bacterial defense systems rapidly diversify to keep pace with counter-evolving phages by swapping effector and sensor components (*43*). The typical modular architecture of defense systems led us to hypothesize that defensive HAM1s might associate with other proteins to form defense systems beyond Kongming. To investigate this, we analyzed the phylogenetic tree of defensive HAM1s and searched for HAM1-associated proteins. Defensive HAM1s can be classified into two major clades and several subclades that have different association protein patterns (Fig. 6E and fig. S19). Defensive HAM1s from all clades are tightly associated with KomC (Fig. 6E and fig. S19), implying that the immune function of HAM1 is likely due to its functional association with SIR2-like effectors. Indeed, we observed a strong co-evolutionary signal between genetically associated KomB and KomC (Fig. 6F), with instances of these domains being fused into a single protein (fig. S19A, candidate type 4). These observations are in line with that the two proteins form a stable complex (Fig. 2B-C). In contrast, neither of these genes showed a comparable correlation with the evolutionary distances of the host 16S rRNA sequences, suggesting that the KomBC module frequently spreads horizontally (Fig. 6F). Together, these results indicate that the KomBC association has remained stable across evolutionary timescales, coupling the immune sensing of second messengers to KomC activation.

By comparison, the association between defensive HAM1 and KomA appears less conserved. In the absence of KomA, we identify various proteins that appear to be genetically associated with the KomBC module (Fig. 6E, fig. S19). Notably, these instances often include proteins involved in nucleotide metabolism or modification (e.g., 5’-nucleotidase, uracil-DNA glycosylase, NTPase, and SAM-dependent methyltransferase), suggesting a conserved function in the generation of dITP or other nucleotide-based immune signaling molecules to activate the KomBC effector (fig. S19A). Interestingly, we found associated RADAR-like systems that may functionally synergize with KomBC by serving as sensor modules, with RdrB harboring an adenosine deaminase domain that is known to drive the massive production of ITP and dITP molecules (fig. S19A) (*44, 45*). Although the ability of these genetically-associated modules to activate KomBC needs further investigation, their common presence in MGEs and within defensive genomic contexts strongly suggests their antiviral function. (fig. S19B). Overall, the diversity of bacterial operons containing KomBC modules and other proteins indicates that various (sub)types of Kongming are widespread across bacteria, reflecting that the modularity of the bacterial immune systems enables their rapid diversification in the arms race with phages.

## Discussion

In this study, we discover and characterize the Kongming system, which mediates a novel bacterial immune signaling pathway based on the production of deoxyinosine 5’-triphosphate (dITP) second messengers. Within the effector complex, a HAM1 domain subunit (KomB) binds the dITP signal and mediates the activation of a SIR2 effector (KomC), which rapidly degrades cellular NAD^+^ to provide population-level immunity (Fig. 5F). Our findings reveal that noncanonical nucleotides serve as a new type of immune second messengers in bacteria.

Nucleotide-based second messengers play vital roles in all three life domains, including response to diverse intracellular stimuli, regulation of basic metabolic adaptations, and immune signaling. Remarkably, prokaryotic anti-phage signaling and eukaryotic innate immune signaling pathways share similar second messengers that robustly activate downstream immune effectors through the amplification of minimal infection cues. These signals include cGMP-AMP (cGAMP), shared by the evolutionary-related cGAS-STING antiviral pathway in animals (*46*) and bacterial CBASS (*10, 11*); cADPR, produced by TIR domains in plants (*7*) and bacteria (*12*); and (cyclic)oligoadenylates, synthesized by oligoadenylate synthetase in animals (*5*) and the type III CRISPR-Cas system in bacteria and archaea (*8, 9*). Beyond the use of cyclic nucleotide signals, it has recently been shown that certain type III CRISPR-Cas systems produce S-adenosyl methionine (SAM)-ATP conjugates (*47*), and a Thoeris type II system rely on histidine-ADPR signaling molecules (*48*), expanding the diversity of immune second messengers to molecules synthesized via conjugation of nucleotides and other moieties. In contrast, Kongming produces signaling molecules via base modification to generate the noncanonical nucleotide dITP, thus representing a new class of nucleotide-based immune second messengers.

The synthesis of the dITP immune signal involves 3 enzymes, including the adenosine deaminase of Kongming (KomA), phage dNMP kinases (DNKs), and a host dNDP kinase (NDK). KomA and phage DNKs can synthesize dIDP from dAMP via two pathways, subsequently followed by NDK conversion of the generated dIDP to dITP (fig. S14F). The two alternative pathways might enable Kongming to defend against a wide range of phages while avoid self-toxicity. First, different phage DNKs may have distinct substrate specificity, probably with weak or little dIMP phosphorylation activity, as dIMP is not a natural substrate for phage dNTP synthesis. Thus, to efficiently trigger immune signaling, KomA shows a relatively high dAMP deamination activity. Notably, dAMP is not the precursor of host dATP synthesis (fig. S6B) and dIMP can be recycled in purine salvage pathway (fig. S6C) (*34*), such that dAMP deamination is not toxic for cells. On the other hand, deamination of host dADP is lethal because the generated dIDP can be rapidly converted to dITP, triggering the immune response. Therefore, the dADP deamination activity of KomA should be weak to avoid self-toxicity. In addition, the host dADP is generated via reduction of ADP by ribonucleotide reductase (RNR)(fig. S6B), the activity of which is tightly regulated (*34*). Such regulation might also protect the host dADP from KomA deamination through an unknown mechanism, so that only the phage-generated dADP serves as the substrate that specifically triggers immune signaling.

We further reveal that T5-like phages encode deoxyribonucleotide 5’ monophosphatase (Dmp) enzymes that counteract Kongming immunity (Fig. 5F). Notably, during the infection cycle of T5, the genomic region encoding Dmp is injected into cells first, resulting in early expression of the enzyme and depletion of cellular dAMP pool (Fig. 5B) (*39, 49*). Afterwards, the rest of the phage genome, including the *dnk* gene, is injected into cells, and phage genome replication is initiated (*39*), with all required nucleotides synthesized de novo (fig. S6D)(*42*). This time-resolved expression of Dmp and DNK may completely avoid dITP synthesis, thereby preventing Kongming activation. In support of this notion, Dmps are primarily distributed in phages that also carry DNKs. Together, these findings reveal a defense and anti-defense arms race for the control of the host’s nucleotide metabolism.

Given that many important components of the eukaryotic innate immune system originate from ancient bacterial immune components (*50*), our findings raise the question as to whether dITP or other non-canonical nucleotides may function as signaling molecules across domains of life. Non-canonical nucleotides are indeed produced in Eukaryotic cells through spontaneous deamination (*51*). Future research directions should investigate whether these non-canonical nucleotides can also trigger immune signaling pathways in eukaryotes. Furthermore, since the KomBC effector complex can be activated by dITP, it provides a potential tool for detecting this non-canonical nucleotide, which can accumulate due to mutation of ITPase and incorporated into DNA during replication, leading to genome instability, cancer, and other diseases (*51, 52*). In conclusion, our discovery of the first non-canonical nucleotide-based anti-phage signaling system expands the universe of nucleotide-based immune secondary messengers, enhancing our understanding of the bacteria-phage arms race, and laying the groundwork for developing future nucleotide detection tools.

## ACKNOWLEDGMENTS

We thank Fangkui Wang at the National Key Laboratory of Agricultural Microbiology, Ting Liu and Linlin Zhong at the National Key Laboratory for Germplasm Innovation & Utilization of Horticultural Crops, Delin Zhang at the core facilities of Center for Protein Research (CPR) and Jinshan Li at Experimental Teaching Center of Bioengineering at Huazhong Agricultural University for technical support. We thank Prof. Shuke Wu for providing Rao1 phage and Dr. Alexander Harms for providing BASEL collection. We thank Prof. Søren J. Sørensen for generously granting us access to the laboratory space and technical equipment at the Section of Microbiology, University of Copenhagen, Denmark. Ruiliang Zhao is a recipient of CSC Scholarship.

## Funding

National Key Research and Development program of China 2022YFA0912200 (W.H.),

National Natural Science Foundation of China 31970545 (W.H.)

National Natural Science Foundation of China 32270099 (W.H.)

Fundamental Research Funds for Central Universities 2662024SKPY003 (W.H.)

Foundation of Hubei Hongshan Laboratory 2021hszd022 (W.H.)

Lundbeck Fonden grant R347-2020-2346 (R.P-R)

VILLUM FONDEN VIL60763 (R.P-R).

Shandong University SKLMT Frontiers and Challenges Project SKLMTFCP-2023-01 (Q.S.)

## Author contributions

Conceptualization: Z.Z., R.P-R, W.H.;

Methodology: Z.Z., R.Z., J.R., H.H;

Investigation: Z.Z., Z.H., R.Z., J.R., M.R.M, Y.L., S.L., H.F., Y.C., N.C., J.Z., D.P., M.L., Q.S.,

Visualization: Z.Z., R.Z.;

Writing – original draft: Z.Z., R.P-R, W.H.;

Writing – review and editing: All authors;

Funding acquisition: Q.S., R.P-R, W.H.;

Supervision: R.P-R, W.H.

## Data and materials availability

All data needed to evaluate the conclusions in the paper are present in the paper and/or the Supplementary Materials. All materials and codes in this paper will be shared upon request. Raw sequencing data are available in Sequence Read Archive (BioProject Accession: PRJNA1149702, PRJNA1195247).

## Supplementary Materials

Materials and Methods

Figs. S1 to S19

References

Table S1 to S7

