## Supplementary materials for "Base-modified nucleotides mediate immune signaling in bacteria"

**The PDF file includes:**

Materials and Methods
Figs. S1 to S19
References

**Other Supplementary Materials for this manuscript include the following:**

Table S1 to S7

### Materials and Methods

#### *Bacterial strains and phages*

*E. coli* DH5α and *E. coli* BL21(DE3) were used for plasmid construction and protein expression, respectively. *E. coli* strains MG1655, PC0886 and ER2738 were used as hosts for phage-related experiments. DH5α and BL21(DE3) were grown in lysogeny broth (LB) medium at 37 °C shaking at 180 rmp. The strains used for phage experiments were grown in MMB medium (LB supplemented with 0.1 mM $MnCl_2$, 5 mM $MgCl_2$) at 37°C shaking at 180 rmp. When appropriate, the media were supplemented with kanamycin (50 μg/ml), streptomycin (50 μg/ml), ampicillin (100 μg/ml) and/or chloramphenicol (25 μg/ml) to maintain the plasmids. All phages used in this study and their respective hosts are listed in table S4. Rao1 phage is a gift from Prof. Shuke Wu at Huazhong Agricultural University and sequenced in this study. Phages of BASEL collection are a gift from Dr. Alexander Harms at University of Basel (*1*).

#### *Plasmid construction*

The plasmids vectors used in this study and the constructed plasmids are listed in table S2. To express the proteins of interest (table S1), their coding sequences, sometimes together with their native promoters as indicated, were amplified from phage or *E. coli* genomic DNA, or synthesized by GENEWIZ (Suzhou, China). In general, the plasmids were constructed with ClonExpress II One Step Cloning Kit (Vazyme, C112-02) or restriction-ligation cloning method as described previously (*2*). Site-directed mutagenesis was performed by PCR amplification of the plasmids carrying the intact genes using the primers containing mutant sites, followed by assembly of the resulting plasmid fragments in *E. coli* DH5α. The primers used in the study are listed in table S3.

The KomABC system, together with its native promoter, was synthesized and inserted into the pET28a plasmid between the BglII and XhoI sites by GENEWIZ (Suzhou, China), resulting in pET28a-KomABC. The plasmids encoding mutated KomABC systems, including pET28a-A$^{m}$HS, pET28a-KomAB$^{m}$C and pET28a-KomABC$^{m}$, were constructed by the site-directed mutagenesis method using primers from ZZF-36 to ZZF-41. The KomABC system with its native promoter and the mutated variants were also cloned into the pCDFDuet-1 vector, generating pCDF-KomABC, pCDF-KomA$^{m}$BC, KomAB$^{m}$C and pCDF-KomABC$^{m}$ respectively.

To express R1DNK, the coding sequence of R1DNK was amplified from Rao1 phage genomic DNA using primers ZZF-5 and ZZF-6, and then inserted into pBAD/His_A plasmid between the NcoI and HindIII sites, generating pBAD-R1DNK. The plasmid encoding R1DNK mutant (pBAD-R1DNK$^{K15A}$) was constructed by the site-directed mutagenesis method using primers ZZF-42 and ZZF-43. The pBAD/His_A-derived plasmids expressing other DNKs were constructed in a similar way. The coding sequences of T5DNK and Escherichia phage KarlJaspers (MZ501090) DNK were amplified from the genomic DNA of the corresponding phages, while the coding sequences of the DNKs from Klebsiella phage vB_KpnS-VAC2 (MZ428221), Bacillus phage vB_BsuM-Goe24 (OM728302), Erwinia phage FIfi106 (OR284297) and Vibrio phage XZ1 (ON000910) were synthesized and inserted into the pBAD/His_A plasmid between NcoI and HindIII sites by GENEWIZ (Suzhou, China).

We constructed a pBAD-Tag vector by inserting the sequence encoding an N-terminal 3× Myc-tag and a C-terminal 3× Flag-tag into pBAD/His_A. The plasmids pBADT-KomA (containing an N-terminal 3× Myc-Tag), pBADT-KomA$^{m}$ (containing an N-terminal 3× Myc-Tag), pBADT-KomB (containing an N-terminal 3× Myc-Tag), pBADT-KomB$^{m}$(containing an N-terminal 3× Myc-Tag), pBADT-KomC (containing an N-terminal 3× Myc-Tag), pBADT-KomC$^{m}$ (containing an N-terminal 3× Myc-Tag), pBADT-R1DNK (containing a C-terminal 3× FLAG-Tag), and pBADT-R1DNK$^{m}$ (containing a C-terminal 3× FLAG-Tag) were constructed for analyzing the expression of mutant proteins. The coding sequence of T5dmp together with its native promoter (without the termination codon) was amplified from T5 phage genomic DNA using primers ZZF-14 and ZZF-15, and then inserted into the pBAD-Tag vector between the SphI and EcoRI sites, resulting in pBADT-T5Dmp. The plasmid encoding T5Dmp mutant (pBADT-T5Dmp$^{D22A}$) was constructed by the site-directed mutagenesis method using primers ZZF-44 and ZZF-45.

To analyze the effects of Dmp on the toxicity triggered by DNK and KomABC, a series of pETDuet-1-derived plasmids that express DNKs or co-express DNKs and Dmps (wild type or mutant) were constructed. At first, we constructed pETDuet-araBAD by replacing a T7 promoter with the araBAD promoter from pBAD/His_A. Then, the coding sequences of different DNKs were inserted into pETDuet-araBAD and their expression is controlled by the araBAD promoter, generating the pETDuet-DNK plasmids. Next, the coding sequences of different Dmps together with the native promoter of T5Dmp were inserted into their respective pETDuet-DNK plasmids, generating the pETDuet-DNK_Dmp plasmids that co-express DNK and Dmp. The pETDuet-T5DNK-T5Dmp$^{D22A}$ was constructed by the site-directed mutagenesis method using primers ZZF-

44 and ZZF-45. The coding sequences of T5DNK and Dmp, Escherichia phage GreteKellenberger DNK and Dmp were amplified from the genomic DNA of the corresponding phages. The coding sequences of Vibrio phage VPG01 DNK and Dmp were synthesized by GENEWIZ (Suzhou, China).

To delete the *dmp* gene of T5, pCas13a-spT5Dmp that targets the *T5dmp* gene and pBAD-ΔT5Dmp that contains two homologous arms flanking the *T5dmp* gene were constructed. For the former, the spacer sequence targeting *T5dmp* was assembled by thermal annealing of two primers ZZF-58 and ZZF-59, and inserted into the linearized pBA559 plasmid (Addgene plasmid # 186235) using the restriction-ligation cloning method. For the latter, the two homologous arms were amplified from the T5 genomic DNA, assembled by overlapping PCR, and inserted into pBAD-Tag between the SphI and EcoRI sites. The plasmids pCas13a-spT5dnk, pCas13a-spR1DNK, pUC19-ΔT5 DNK, and pUC19-ΔR1DNK were constructed by the same method as described above for deleting the *dnk* genes of T5 and Rao1.

pQLink-R1DNK (containing an N-terminal Strep-Tag II), pQLink-T5Dmp (containing an N-terminal Strep-Tag II), pQLink-T5Dmp$^{D22A}$ (containing an N-terminal Strep-Tag II), pCDF-KomC (containing an N-terminal 6×His tag), pCDF-KomA (containing an N-terminal 6×His tag), pQLink-NDK (containing an N-terminal 6×His tag), and pMAL-KomA (containing an N-terminal MBP tag) were constructed to express the recombinant proteins in *E. coli* BL21(DE3) for protein purification. The coding sequence of NDK was amplified from E. coli MG1655 genomic DNA, while the sequences of other genes were amplified from the previously constructed plasmids respectively. The sequences were inserted into the indicated vectors using ClonExpress assembly. To co-express R1DNK and KomA, the R1DNK expression cassette (containing the araBAD promoter) was amplified from the pBAD-R1DNK plasmid and then cloned into pRSFDuet-1, generating pRSF-R1DNK. Next, the coding sequence of KomA was amplified from pET28a-KomABC plasmid and inserted in the second multiple cloning site of pRSF-R1DNK, generating pRSF-R1DNK-KomA.

To co-express His-tagged KomC and KomB, the coding sequence of KomB was amplified from pET28a-KomABC plasmid using primers ZZF-24 and ZZF-25, and inserted into the second multiple cloning site of pCDF-KomC, generating pCDF-KomC-KomB. To co-express untagged KomC and KomB, PCR amplification was performed using pCDF-KomC-KomB as template and the primers ZZF-60 and ZZF-61. The resulting plasmid fragment was assembled using ClonExpress assembly (Vazyme, Nanjing, China), generating pCDF-KomC(tag-free)-KomB. The plasmids expressing the indicated mutated proteins were constructed by the site-directed mutagenesis method using the primers as listed in table S3.

#### *Protein expression and purification*

To obtain the recombinant proteins, the corresponding plasmids were transformed into *E. coli* BL21(DE3). The transformants were grown to an optical density ($OD_{600}$) of ~1.0 at 37 $^{o}$C in LB medium supplemented with appropriate antibiotics. Then, protein expression was induced with 0.4 mM isopropyl-β-D-thiogalactoside (IPTG) and/or 0.2% L-arabinose overnight at 16 $^{o}$C, except for T5Dmp and T5Dmp$^{D22A}$, the expression of which was induced with 0.4 mM IPTG at 37 ℃ for only 5 min. The cells were collected by centrifugation at 7000 g for 10 min and the cell pellets were stored in -80 $^{o}$C freezer before protein purification.

To purify the His-tagged proteins (KomB-KomC complex, KomC, KomA, NDK), cell pellets were resuspended in lysis buffer (20 mM HEPES pH 7.5, 20 mM imidazole, 250 mM NaCl) and lysed by French press. The lysates were centrifuged at 13000 g for 1 h at 4 $^{o}$C to remove cell debris, and the supernatants were then loaded onto a Ni-NTA column (Cytiva, 17531801) pre-equilibrated with lysis buffer. The column was washed with wash buffer (20 mM HEPES pH 7.5, 60 mM imidazole, 250 mM NaCl) and then the His-tagged proteins were eluted with elution buffer (20 mM HEPES pH 7.5, 300 mM imidazole, 250 mM NaCl). The eluted proteins were concentrated with Amicon centrifuge filters (Millipore, UFC8010) and then loaded onto a Superdex 200 Increase 10/300 GL column (Cytiva, 28990944) pre-equilibrated with buffer C (20 mM Tris-HCl, pH 7.5, 250 mM NaCl). The eluted proteins were concentrated again and stored at -80 $^{o}$C before use. The purity was analyzed by SDS-PAGE.

To purify the Strep-tagged protein (R1DNK, T5Dmp and T5Dmp$^{D22A}$), the cell pellet was resuspended in buffer C (20 mM Tris-HCl, pH 7.5, 250 mM NaCl) and lysed by French press. The lysate was centrifuged at 13000 g for 1 h at 4 $^{o}$C to remove cell debris, and the supernatant was then loaded onto a Strep-Tactin®XT column (IBA Life Sciences, 2-5030-025) pre-equilibrated with buffer C. The column was washed with buffer C and then the Strep-tagged protein was eluted with buffer BXT (20 mM Tris-HCl, pH 7.5, 250 mM NaCl, 50 mM biotin). The eluted protein was concentrated with an Amicon centrifuge filter (Millipore, UFC8010) and then loaded onto a 5 ml HiTrap Desalting column (Cytiva, 29048684) pre-equilibrated with buffer C (20 mM Tris-HCl, pH 7.5, 250 mM NaCl). The eluted proteins were concentrated again and stored at -80 $^{o}$C before use. The purity was analyzed by SDS-PAGE.

Protein co-purification was performed to analyze the physical interaction between KomC and KomA, KomB or R1DNK. Specifically, *E. coli* BL21(DE3) cells carrying pRSF-R1DNK-KomA and pCDF-KomC$^{m}$-KomB (His-tagged KomC) were grown to an $OD_{600}$ ~1.0 at 37 °C in LB medium supplemented with appropriate antibiotics. Protein expression was induced with 0.4 mM IPTG and 0.2% L-arabinose at 16 °C for 16 h. The cell lysate preparation and Ni-NTA affinity chromatography were carried out using the same method for His-tagged proteins as described above. In addition, protein expression and purification were also performed with *E. coli* BL21(DE3) cells carrying pRSF-R1DNK-KomA and pCDF-KomC$^{m}$ (tag-free)-KomB (untagged KomC). The fractions from the Ni-NTA affinity chromatography were subjected to mass spectrometry.

To obtain the putative second messenger bound by the KomB-KomC complex, *E. coli* BL21(DE3) cells co-expressing R1DNK or its mutant and KomABC or the mutated systems were used to purify the different versions of KomBC complexes, i.e. KomBC-1 (pCDF-KomC-KomB and pRSF-R1DNK$^{K15A}$-KomA), KomBC-2 (pCDF-KomC-KomB and pRSF-R1DNK-KomA$^{m}$), H$^{m}$S (pCDF-KomC-KomB$^{m}$ and pRSF-R1DNK-KomA) and KomBC$^{m}$ (pCDF-KomC$^{m}$-KomB and pRSF-R1DNK-KomA) (Fig. 3C). Culture growth, protein expression induction and Ni-NTA affinity chromatography were carried out with the same method as described above, except the column was washed with lysis buffer instead of the wash buffer. The elution fractions were concentrated to 1 ml using Amicon centrifuge filters (Millipore, UFC8010) and analyzed by SDS-PAGE (fig. S11A). The concentrated protein samples were denatured to release the bound molecules.

##### *Phage propagation and plaque assay*

Phages were propagated using *E. coli* MG1655 as the host. Overnight cultures of *E. coli* MG1655 were diluted 1:100 in MMB medium (LB supplemented with 0.1 mM $MnCl_2$, 5 mM $MgCl_2$) and grown to an $OD_{600}$ of ~0.3 at 37 °C. Then, 1 ml of the culture was infected with phages of interest at a multiplicity of infection (MOI) of 0.01 and further grown for 4 h. The cultures were centrifuged at 5000 g for 10 min and the supernatants were filtered through 0.22 µm membrane filters (Millipore, SLGPM33RS) to remove the cells. The resulting phage shocks were stored at 4 °C.

A double agar overlay assay was used to determine phage titre. In brief, phage stocks were 10-fold serially diluted in SM buffer (50 mM Tris-HCl, pH 7.5, 100 mM NaCl, 8 mM $MgSO_4$). Three µl of the serial dilutions (from $10^{-3}$ to $10^{-8}$) were mixed with 300 µl of the overnight cultures of the *E. coli* host as indicated before and 4 ml melted MMB + 0.5% agar, and then poured onto MMB + 1.5% agar plates. Plates were incubated at 37 °C overnight. Plaque formation units (PFUs) were determined by counting the derived plaques.

For plaque assays, *E. coli* host cells expressing an empty vector (pET28a) or wild type or mutated KomABC systems were grown in MMB medium supplemented with kanamycin (50 µg/ml) at 37 °C overnight. Then, 300 µl of the cultures were mixed with 8 ml melted MMB + 0.5% agar + kanamycin (50 µg/ml) and poured onto MMB + 1.5% agar + kanamycin (50 µg/ml) plates. Meanwhile, phages of interest were 10-fold serially diluted in SM buffer and 3 µl of the 10-fold serial dilutions were spotted on the bacterial overlay. The plates were incubated at 37 °C for 8 h before images of the plates were captured.

To analyze the function of T5Dmp, The *E. coli* MG1655 cells carrying pCDFduet empty vector or pCDF-KomABC were further transformed with an empty vector (pBAD/His_A) or pBAD-T5Dmp or pBAD-T5Dmp$^{D22A}$. The transformants were grown in MMB medium supplemented with ampicillin (100 µg/ml) and kanamycin (50 µg/ml) at 37 °C overnight. Then, 300 µl of cultures were mixed with 8 ml melted MMB + 0.5% agar + ampicillin (100 µg/ml) and kanamycin (50 µg/ml) and poured onto MMB + 1.5% agar + ampicillin (100 µg/ml) and kanamycin (50 µg/ml) plates. Meanwhile, phage Rao1 was 10-fold serially diluted in SM buffer and 3 µl of the 10-fold serial dilutions were spotted on the bacterial overlay. The plates were incubated at 37 °C for 8 h before images of the plates were captured.

**Phage-infection dynamics in liquid medium**

Overnight cultures of *E. coli* MG1655 containing an empty vector (pET28a) or pET28a-KomABC were diluted 1:100 in MMB medium supplemented with kanamycin (50 mg/ml) and grown to $OD_{600}$~0.2 at 37 °C. Then, 180 µl of the cultures were transferred into wells of a 96-well plate, and 20 µl of the diluted phages were added into the cultures for a final MOI of 0.3 or 3. The uninfected culture (MOI = 0) was supplemented with 20 µl SM buffer. $OD_{600}$ was measured every 5 min with shaking at 200 rpm at 37 °C in a FLUOstar OMEGA (BMG Labtech, Germany).

##### *Toxicity assays on solid medium*

To analyze the toxicity triggered by the co-expression of the KomABC system and R1DNK, the colony-forming units (CFUs) per ml of the cells co-expressing KomABC and R1DNK were determined. Specifically, *E. coli* MG1655 cells containing pCDFduet-derived plasmids (pCDFduet empty vector or pCDF-KomABC) and pBAD/His_A -derived plasmids (pBAD/His_A empty vector, pBAD-R1DNK or pBAD-R1DNK$^{K15A}$) were grown overnight at 37 °C in MMB supplemented with streptomycin (50 µg/ml) and ampicillin (100 µg/ml). The overnight cultures were then diluted 1:100 in MMB supplemented with appropriate antibiotics and grown to $OD_{600}$~0.8 at 37 °C. The cultures were subsequently 10-fold serially diluted in MMB. Five µl of 10-fold serial dilutions (from $10^{-1}$-$10^{-6}$) were spotted on MMB plates supplemented with appropriate antibiotics and 1% glucose or 0.05% L-arabinose. The plates were incubated at 37 °C overnight before images of the plates were captured. The toxicity triggered by the co-expression of the KomABC system and DNKs from other phages was analyzed in a similar procedure.

To analyze the effects of T5Dmp on the toxicity triggered by the co-expression of the KomABC system and T5DNK, The CFUs per ml of *E. coli* MG1655 cells containing pCDFduet-derived plasmids (pCDFduet empty vector or pCDFduet-KomABC) and pETduet-derived plasmids (pETduet-T5DNK, pETduet-T5DNK-T5Dmp or pETduet-T5DNK-T5Dmp$^{D22A}$) were determined as described above. The effects of Dmps from other phages on the DNK-triggered toxicity were analyzed in the same protocol.

**Toxicity assays in liquid medium**

Single colonies of *E. coli* MG1655 harboring pCDFduet empty vector and pBAD-R1DNK, pCDF-KomABC and pBAD-R1DNK, or pCDF-KomABC and pBAD-R1DNK$^{K15A}$ were grown at 37 $^{o}$C overnight in MMB supplemented with streptomycin (50 µg/ml), ampicillin (100 µg/ml). Overnight cultures were diluted 1:100 in MMB supplemented with streptomycin (50 µg/ml), ampicillin (100 µg/ml) and 0.2% arabinose, and 200 µl of diluted cultures were transferred into wells of a 96-well plate. The plate was incubated at 37 $^{o}$C with shaking at 200 rpm in a FLUOstar OMEGA (BMG Labtech, Germany) and $OD_{600}$ was measured every 5 min.

#### *Isolation of phage escape mutants*

The plaque assay was used to isolate mutant phages that evaded the defense of the KomABC system, as described previously (*3*). Briefly, 300 µl of overnight culture of *E.coli* MG1655 harboring pET28a-KomABC plasmid was mixed with 8 ml melted MMB + 0.5% agar + kanamycin (50 µg/ml) and poured onto MMB + 1.5% agar + kanamycin (50 µg/ml) plates. Three µl of the 10-fold serially diluted Rao1 phage stock was spotted on the bacterial overlay, and then the plates were incubated at 37 $^{o}$C overnight. Single plaques were isolated and propagated using *E. coli* MG1655 at 37 $^{o}$C for 4 h. The cultures were centrifuged for 10 min at 5000 g and the supernatants were filtered through 0.22 µm membrane filters to remove the cells. The resulting phage solution of Rao1 phage mutants was 10-fold serially diluted in SM buffer. Three µl of the 10-fold serial dilutions of wild type Rao1 and Rao1 mutants were spotted on the bacterial overlay of *E. coli* MG1655 containing pET28a or pET28a-KomABC. The plates were incubated at 37 $^{o}$C for 8 h and then images of the plates were captured.

#### *Extraction and sequencing of phage DNA*

To extract phage DNA, 500 µl of phage stocks were treated with DNase I (5 U/ml) (Thermo Fisher Scientific, EN0521) and RNase A (50 µg/ml) (Thermo Fisher Scientific, R1253) at 37 $^{o}$C for 1 h to degrade bacterial DNA and RNA. The phage stocks were then incubated with proteinase K (0.5 mg/ml) (Thermo Fisher Scientific, EO0491) at 37 $^{o}$C for 1 h, followed by supplementation of 500 µl of phenol/chloroform/isoamyl alcohol (pH 8.0, 25:24:1). The samples were then vortexed briefly and centrifuged at 13000 g at 4 $^{o}$C for 15 min. The upper water phase (about 400 µl) was transferred into a 1.5 ml tube and mixed with 400 µl of chloroform. The mixtures were then vortexed briefly and centrifuged at 13000 g for 15 min at 4 $^{o}$C. The upper water phase (about 400 µl) was isolated to a 1.5 ml tube and mixed with 40 µl 3 M NaAc (pH 5.2) and 400 µl isopropanol. The mixtures were then vortexed briefly and incubated at -20 $^{o}$C for 4 h. After incubated, the mixtures were centrifuged at 13000 g at 4 $^{o}$C for 15 min and the supernatant was discarded. The pellets were washed with 75% ethanol (pre-cooled to -20 $^{o}$C) and dried for 10 min. The pellets were resuspended in 30 µl DEPC $H_2O$.

Libraries preparation and sequencing were performed by Novogene (Beijing, China). Briefly, sequencing libraries were prepared using Annoroad® Universal DNA Fragmentase kit V2.0 (AN200101-L) and Annoroad® Universal DNA Library Prep Kit V2.0 (AN200101-L) according to the instructions of the manufacturer. Libraries were sequenced on Novaseq 6000 S4 platform with PE150 strategy.

The sequencing data with the host genome removed were aligned to phages using bwa (v0.7.17-r1188) (*4*). Duplicate reads in the bam files were removed using picard MarkDuplicates (v3.1.1). Variant detection was conducted using bcftools (v1.18) (*5*), starting with variant calling using mpileup (option "--output-type u") and call (option "--multiallelic-caller --variants-only --output-type v"), followed by filtering with filter (option "-g 3 -G 10 -e 'QUAL<40 || DP<10' "). Finally, SNPs (single nucleotide polymorphisms) and indels (insertion-deletions) were detected with bcftools view. The annotation of variant results was performed using the annovar software (option "-geneanno --neargene 100").

**Quantification of relative cellular $NAD^+$ levels**

The relative content of cellular $NAD^+$ was measured by the MTT (Methyl Thiazolyl Tetrazolium) assay as described previously (*2*). Specifically, *E. coli* MG1655 cells expressing wild-type or mutant KomABC and wild-type or mutant R1DNK were grown at 37 $^{o}$C overnight in MMB supplemented with streptomycin (50 µg/ml), ampicillin (100 µg/ml). Overnight cultures were diluted 1:100 in MMB supplemented with appropriate antibiotics and grown at 37 $^{o}$C until $OD_{600}$

reached 0.2. Then, expression of R1DNK or its mutant were induced with 0.2% L-arabinose for 30 min at 37 °C. 4 ml of cultures ($OD_{600}$ ~0.3) were collected by centrifugation at 4000 g for 5 min. The supernatant was removed and the cell pellets were washed with 600 µl PBS buffer (pH 7.4). The cells were collected again by centrifugation at 4000 g for 5 min again. The cellular $NAD^+$ content was measured using the Coenzyme I NAD(H) content detection kit (Solarbio, BC0315) according to the instructions of the manufacturer. The optical density at 570 nm of each sample was measured to reflect the $NAD^+$ content in the sample. The relative $NAD^+$ content of each sample was calculated using the following equation:

$$\text{Relative } NAD^+ \text{ content} = (OD^{sample} - OD^{sample\_control}) / (OD^{EV} - OD^{EV\_control})$$

Control, the $NAD^+$ in the sample was neutralized before supplementation of MTT; EV, the cells containing empty vectors. All samples were performed in three biological replicates.

**Pull-down assay**

To analyze the physical interaction between R1DNK and KomA, KomB-KomC complex (KomBC), the bait protein (20 µg Strep-R1DNK) was incubated with 20 µg purified KomA or 20 µg purified KomBC, or a mixture of them in 200 µl buffer C at 4 °C for 30 min. Then, the protein mixtures were incubated with pre-equilibrated 40 µl Strep-Tactin resin (IBA Life Sciences, 2-5030-025) at 4 °C for 30 min. The resins were washed with 500 µl buffer C for five times. The bound protein was eluted in 100 µl BXT buffer and analyzed by SDS-PAGE.

To analyze the physical interaction between KomA and KomBC, the bait protein (20 µg MBP or MBP-KomA) was incubated with 20 µg purified KomBC in 200 µl buffer C at 4 °C for 30 min. Then, the protein mixtures were incubated with pre-equilibrated 80 µl Dextrin Beads (Smart-Life sciences, SA026005) at 4 °C for 30 min. The Beads were washed with 500 µl buffer C for three times. The bound protein was eluted in 100 µl buffer C supplemented with 10 mM maltose and analyzed by SDS-PAGE.

**Western blot**

For protein mutant expression analysis, *E. coli* MG1655 strain harboring the corresponding plasmid was grown to an $OD_{600}$ of ~0.3 at 37 °C in MMB medium supplemented with appropriate antibiotics and 0.2% arabinose. 5 ml of cultures were collected and resuspended in 600 µl buffer C. Cells were disrupted by ultrasonication, followed by centrifugation at 13000 g at 4 °C for 15 min to remove the debris. The proteins were separated by SDS-PAGE and then transferred to PVDF membranes (Bio-Rad, Hercules, CA, USA). Membranes were blocked in TBST buffer (50 mM Tris, 100 mM Nacl, 0.05% Tween 40, pH 8.0) supplemented with 6% milk at 25 °C for 2 h, followed by incubation with anti-FLAG (Abbkine, ABT2011,1:4000) or anti-Myc (Abbkine, ABT2061, 1:5000) primary antibodies diluted in TBST buffer supplemented with 6% milk overnight at 4 °C. Membranes were washed three times with TBST buffer, followed by incubation with secondary antibodies (Abbkine, A21020, 1:10000) at 25 °C for 2 h and visualized using Tanon 5200 (Tanon, Shanghai, China).

**Deoxynucleotide monophosphate kinase assay**

For deoxynucleotide monophosphate kinase assay, a 50-µl reaction mixture containing 50 mM HEPES pH 7.5, 50 mM KCl, 10 mM $MgCl_2$, 1 mM ATP, 1 mM TCEP, 1 mM nucleotide substrate (dAMP, dGMP, dCMP, dTMP, dIMP, AMP, or IMP) and 1 µM R1DNK was incubated at 37 °C for 1 h. Then, the mixture was heated to 90 °C for 5 min to stop the reactions, followed by centrifugation at 13000 g at 4 °C for 15 min to remove the precipitated protein. The supernatant was supplemented with 750 µl of buffer A (5 mM Tris-HCl pH 8.0) and loaded onto a MonoQ 5/50 column (GE Healthcare, USA). The bound nucleotides were eluted using a linear gradient from 0 to 0.4 M NaCl over 25 ml.

The nucleotides used for all experiments were purchased from the following companies. dGMP (A600363), dCMP (A600361), dTMP (A600367), dUTP (B600006) and 2'-deoxyadenosine (A620043) were purchased from Sangon Biotech. dAMP (S18087), dIMP (S18105), AMP (B24464), IMP (S18082), ADP (B25056), dATP (S18090), dCDP (S18096), dTDP (S18109), inosine (B20582) and ITP (S18084) were purchased from Yuanye. ATP ( R0441) and dITP (R1191) were purchased from Thermo Fisher Scientific. dADP (GC11854) was purchased from GlpBio. 2'-deoxyinosine (532038) was purchased from J&K Scientific. dGDP (D119530) was purchased from Aladdin.

**Adenosine deaminase assay**

To perform the adenosine deamination assay, a 50-µl reaction mixture containing 50 mM HEPES pH 7.5, 50 mM KCl, 1 mM TCEP, 1 mM substrate (dATP, dADP, dAMP, ATP, ADP or AMP) and 5 µM KomA was incubated at 37 °C for 1 h. After heated to 90 °C for 5 min to stop the reactions, the reaction mixtures were supplemented with 1.5 µl FastAP Thermosensitive Alkaline Phosphatase (Thermo Fisher Scientific, EF0654) and incubated at 37 °C for 15 min. Then, the mixtures were supplemented with 350 µl MilliQ $H_2O$ and centrifuged at 13000 g for 10 min to remove precipitated protein. The supernatant was passed through 0.2 µm Filters for Ultra high-performance liquid chromatography (UHPLC) analysis.

The HPLC analysis was performed using a Nexera UHPLC System (Shimadzu, Japan) with a C18 column (Kinetex 3×100 mm, particle size 2.6 µm) for both the reaction products and the standards. The column was warmed to 35 °C and the flow rate was 0.3 ml/min. Three µl of each sample was loaded. Gradient elution was performed with solvent A (5 mM ammonium acetate) and solvent B (100% acetonitrile). The gradient elution proceeded as follows: 0 to 2 min, 5% B; 2 to 7 min, 5 to 10% B; 7 to 10 min, 10 to 95% B; 10 to 18 min, 5% B.

To analyze the effects of divalent metal ions on the adenosine deamination activity of KomA, the assay was performed with dAMP as substrate in the presence of 0.5 mM different divalent metal ions $Mg^{2+}$, $Ni^{2+}$, $Cu^{2+}$, $Zn^{2+}$, $Mn^{2+}$) or 0.5 mM EDTA.

**εNAD-based NADase assay**

To analyze the NADase activity of KomB-KomC complex (KomBC) and the mutated complexes, a 50-µl reaction mixture containing 50 mM HEPES pH 7.5, 50 mM KCl, 1 mM TCEP, 50 µM nicotinamide 1, $N^6$-ethenoadenine dinucleotide (εNAD, Sigma, N2630), and the indicated enzymes, nucleotides, metal ions and/or reaction products was prepared in black 96-well half area plates (Greiner Bio-One, 675076). Then, the plates were immediately incubated in a Spark multimode microplate reader (Tecan, Switzerland) at 37 $^oC$ and fluorescence intensity was measured every 30 s over 30 min at 300 nm excitation wavelength and 410 nm emission wavelength. The concentrations of the KomBC complex in the reaction were calculated based on the estimation that the ratio of KomB and KomC is 1:1 according to the band density in SDS-PAGE gel and the protein mass spectrometry analysis.

To analyze the NADase activity of standalone KomC and KomBC complex (Fig.2E), the reaction mixtures contained 0.1 mM $ZnSO_4$ and 0.8 µM KomC or KomBC.

To analyze the effects of divalent metal ions on the NADase activity (fig. S10A), the reaction mixtures contained 0.8 µM KomBC and 1 mM divalent metal ions ($Mg^{2+}$, $Ni^{2+}$, $Cu^{2+}$, $Zn^{2+}$, $Mn^{2+}$) or 1 mM EDTA. Except for the metal dependency assay, all other reactions contained 0.1 mM $ZnSO_4$.

To analyze whether the NADase activity is activated by nucleotides (Fig. 3A), the reaction mixtures contained 50 nM KomBC and 100 nM different nucleotides (dNTP, NTP, dITP, dUTP, ITP, IDP, IMP). The effects of dITP and IDP on the NADase activity were further analyzed with a gradient of dITP (0, 0.1 nM, 1 nM, 10 nM, 100 nM) and IDP (0, 10 nM, 100 nM, 1000 nM) (fig. S10C-D).

To analyze the NADase activity of mutated KomBC complexes ($KomB^m$-KomC and KomB-$KomC^m$), the concentrations of wild type KomBC, $KomB^m$-KomC or KomB-$KomC^m$ in the reaction mixtures were 0.6 µM (Fig. 2G). To analyze the activation of mutated KomBC complexes by dITP, the concentrations of wild type and mutated complexes were 50 nM and 1 nM dITP was further supplemented (Fig. 3B).

To analyze whether the reaction products of KomA and *E. coli* NDK with dADP as substrate and ATP as phosphate group donor activate the NADase activity of KomBC, the adenosine deamination reaction mixtures containing 50 mM HEPES pH 7.5, 50 mM KCl, 1 mM TCEP, KomA (0 or 5 µM), 100 nM dADP, were incubated at 37 $^oC$ for 60 min, and further supplemented with 100 µM $ZnSO_4$,10 mM $MgCl_2$, NDK (0 and 1.6 µM) and 1 µM ATP, followed by incubation at 37 $^oC$ for 15 min. Then, the mixtures were heated at 90 $^oC$ for 5 min to stop the reactions. After the precipitated protein was removed by centrifugation at 13000 g at 4 $^oC$ for 15 min, the reaction products were tested for its capability of activating the NADase activity of KomBC in reaction mixtures containing 50 nM KomBC (Fig. 4F).

To analyze whether the reaction products of R1DNK and *E. coli* NDK with dIMP as substrate and ATP as phosphate group donor activate the NADase activity of KomBC, the reaction mixtures containing 50 mM HEPES pH 7.5, 50 mM KCl, 1 mM TCEP, 10 mM $MgCl_2$, R1DNK (0 or 2.4 µM), 1 mM ATP, 1 µM dIMP, were incubated at 37 $^oC$ for 60 min, and further supplemented with 100 µM $ZnSO_4$ and NDK (0 or 1.6 µM), followed by incubation at 37 $^oC$ for 15 min. Then, the mixtures were heated at 90 $^oC$ for 5 min to stop the reactions. After the precipitated protein was removed by centrifugation at 13000 g at 4 $^oC$ for 15 min, the reaction products were tested for its capability of activating the NADase activity of KomBC in reaction mixtures containing 50 nM KomBC (Fig. 4G).

To analyze whether the released molecules from the denatured protein samples (KomBC-1, KomBC-2, $KomB^mC$ and $KomBC^m$) activate the NADase activity of KomBC, 1 ml of KomBC-1, KomBC-2, $KomB^mC$ and $KomBC^m$ were heated to 90 $^oC$ for 10 min and then centrifuged at 13000 g at 4 $^oC$ for 15 min to remove the precipitated protein. Then, NADase assay were performed in a 50-µl reaction mixture containing 50 nM KomBC and 0.01% (v/v) of the heated samples of KomBC-1, KomBC-2, $KomB^mC$ and $KomBC^m$ (fig. S11C).

**UHPLC analysis of the NADase reaction products**

To analyze the products of NADase reactions, a 50-µl reaction mixture containing 50 mM HEPES pH 7.5, 50 mM KCl, 1 mM TCEP, 1 mM $NAD^+$ (Yuanye, B27081), 100 µM $ZnSO_4$, 100 nM KomBC and dITP (0 or 1 nM) was incubated at 37 $^oC$ for 30 min. The reaction mixtures were heated to 90 $^oC$ for 5 min and then 450 µl MilliQ $H_2O$ was supplemented. The mixtures were then centrifuged at 13000 g for 10 min to remove precipitated protein and the supernatants was passed through 0.2 μm Filters for UHPLC analysis.

The reaction products and standards ($NAD^+$(Yuanye, B27081), ADPR (MCE, HY-100973A), NAM (Yuanye, B21424)) were analyzed using a Nexera UHPLC System (Shimadzu, Japan) with a C18 column (Kinetex, 3×100 mm, particle size 2.6 μm). The column was warmed to 30 $^{o}$C and the flow rate was 0.3 ml/min. Three μl of each sample was loaded. Gradient elution was performed with solvent A (0.1% (v/v) formic acid and 5 mM ammonium formate in water) and solvent B (0.1% (v/v) formic acid and 5mM ammonium formate in methanol). The gradient elution proceeded as follows: 0 to 5 min, 0% B; 5 to 7 min, 0% to 95% B; 7 to 7.5 min, 95% B; 7.5 to 8 min, 95% to 0% B, 8 to 21 min 0% B.

**Phage T5 genome editing**

T5 *dmp* gene was knocked out following a published method (*6*) with modifications. Briefly, *E. coli* MG1655 cells harboring pBAD-ΔT5dmp plasmid (a homologous recombination plasmid) were grown in MMB medium with ampicillin (100 μg/ml) overnight at 37 $^{o}$C. The overnight culture was diluted in MMB medium with ampicillin (100 μg/ml) to $OD_{600}$~0.2. Then, 500 μl of the culture was infected with wild type T5 phage at an MOI of 0.01 and grown at 37 $^{o}$C for 4 h. The culture was centrifuged for 10 min at 5000 g and the supernatants were filtered through 0.22 μm membrane filters to remove the cells. This resulted in a mixture of wild type T5 phage and homologous recombination-edited T5 phage. The phage titre of the mixture was determined using the double agar overlay assay as described above.

Then, *E. coli* MG1655 cells harboring the pCas13a-spT5dmp plasmid, which carried a spacer targeting *dmp*, were grown in MMB medium containing chloramphenicol (25 μg/ml) overnight at 37 $^{o}$C. The overnight culture was diluted in MMB medium supplemented with chloramphenicol (25 μg/ml) and 80 ng/ml aTc to $OD_{600}$~0.2. Then, 500 μl of the culture was infected with the mixture of wild type T5 phage and homologous recombination-edited T5 phage at an MOI of 0.01 and grown at 37 $^{o}$C for 4 h. The culture was centrifuged for 10 min at 5000 g and the supernatant was filtered through 0.22 μm membrane filters to remove the cells. The resulting phage mixture was used to infect the cells harboring the pCas13a-spT5dmp plasmid again to enrich the homologous recombination-edited T5 phage.

To select the homologous recombination-edited T5 phage, the mixture of wild type T5 phage and homologous recombination-edited T5 phage were 10-fold serially diluted in SM buffer and 3 μl of 10-fold serial dilutions were spotted on the bacterial overlay prepared with *E. coli* MG1655 cells harboring pCas13a-spT5dmp. Then, the plates were incubated at 37 $^{o}$C for 8 h and single plaques were analyzed with PCR using the primer set ZZF-14 and ZZF-15. The propagation of T5 mutants was performed with *E. coli* MG1655 cells as the host as described above.

Similar method was used to knockout T5DNK and R1DNK.

**Fitness assay of mutant phages**

Phages competition assays were performed following a published method (*3*) with minor modifications. Each pair of mutant and wild-type (WT) phages was mixed at an approximate ratio of 1:1 as the ancestor of phage mixture (cycle 0). The cultures of *E. coli* MG1655 were grown to an $OD_{600}$ of 0.2 at 37 $^{o}$C shaking at 180 rmp. Then the cultures were infected with the phage mixture at a final MOI ~ 0.01, and incubated for 4 h at 37 $^{o}$C shaking at 180 rmp. the culture was centrifuged at 8000 g for 5 min to remove cells and debris, and the phage titer in the supernatant was assessed using the same method as described above. Subsequently, a fresh culture of *E. coli* MG1655 were infected with the resulting supernatant at a final MOI ~ 0.01 (cycle 2). These cycles (1, 2, and 3) were performed three times, and these resulting phage mixtures were sampled for phage DNA Extraction and sequencing as described above. For the origin of the phage mixture of Rao1 and $Rao1^{\Delta dnk}$ (cycle 0), 500 μl of the phage mixture was extracted for sequencing.

The sequenced reads were aligned with the mutant phage genome sequence, and the ratio of each mutant reads to WT reads was calculated based on the reads aligned at the expected mutation position. The sequencing reads were aligned to phages genome using bwa (v0.7.17-r1188).

**dITP pyrophosphatase assay of KomBC**

To analyze whether KomBC has dITP pyrophosphatase activity, a-50 μl reaction mixture containing 50 mM HEPES pH 7.5, 50 mM KCl, 1 mM TCEP, 100 μM $ZnSO_4$, KomBC (0 or 3.2 μM) and 1 mM dITP was prepared and incubated at 37 $^{o}$C for 60 min. The reaction mixtures were heated to 90 $^{o}$C for 5 min to stop the reaction, and the reaction products were analyzed using ion-exchange chromatography as described in the **Deoxynucleotide monophosphate kinase assay**.

**Deoxynucleoside 5'-phosphatase assays**

Deoxynucleoside 5'-phosphatase activity of T5Dmp was analyzed following a published method (*7*) with modifications. Briefly, the reactions were prepared in a 50 μl reaction mixture containing 50 mM Tris-HCl pH 7.5, 50 mM NaCl, 0.1 mg/ml BSA, 4 mM DTT, 30 mM $MgCl_2$, 0.5 μM T5Dmp protein (wild type or mutant) and 1 mM different deoxynucleotide substrates (dNMP, dITP, dATP, dADP, dIMP) at 30 °C for 30 min. The reaction mixtures were heated to 90 °C for 5 min to end reactions. The reaction products were analyzed using UHPLC as described in the Deaminase assay.

**Protein identification by mass spectrometry (MS)**

Protein MS was performed by Novogene (Beijing, China). To identify the purified protein bands, i.e. the putative KomB protein purified from cells co-expressing His-KomC and KomB, the protein samples were separated by 10% SDS-PAGE and visualized with Coomassie brilliant blue. The protein band indicated in Fig. 2C was excised and sent for MS analysis by Novogene (Beijing, China). Briefly, the proteins in gel were digested by trypsin with a standard protocol of Novogene, and the digested peptides were analyzed by an EASY-nLC™ 1200 UHPLC (Thermo Fisher) coupled with an Q Exactive™ HF-X mass spectrometer (Thermo Fisher) using a standard LC-MS/MS method.

MS was also used to identify the proteins co-purified with His-tagged KomC or untagged KomC (Fig. 2B). The proteins were also digested by trypsin. The digested peptides were desalted with a C18 desalting column and lyophilized. Then, the digested peptides were analyzed by a nanoElute UHPLC (Bruker) coupled with a tims TOF pro2 (Bruker) using a standard LC-MS/MS method.

Peptide spectrum matches (PSMs) were performed in Proteome Discoverer (PD, Thermo) or MaxQuant (Bruker, Tims) against *Escherichia coli* (strains B / BL21-DE3) proteome database (Uniprot, UP000009074) and the 4 additional proteins (KomA, KomB, KomC, R1DNK) with their amino acid sequences. The search results were filtered with PD/MaxQuant software to improve the quality of the search results. PSMs with a confidence level of 99% or more were considered as credible PSMs. The identified proteins contain at least 1 unique peptide. False discovery rate (FDR) validation was performed to remove the peptides and proteins with FDR greater than 1%. To determine the significance of the differences, the Significance A test was performed on the relative quantitative values of each protein in the two comparison sample pairs (*8*), and the corresponding P-value was calculated. Proteins with significant quantitative differences between the His-tagged KomC group and the untagged KomC group ($p < 0.1$, $|\log_2 FC| > 1$ ($FC > 2$ or $FC < 0.5$ [fold change, FC]) were defined as differentially expressed proteins (DEP).

**Liquid chromatography-tandem mass spectrometry (LC-MS/MS)**

LC-MS/MS was used to identify the released molecules from the denatured protein samples (KomBC-1, KomBC-2, $KomB^mC$ and $KomBC^m$) and the reaction products of KomA and *E. coli* NDK with dADP and ATP. For the former, 1 ml of KomBC-1, KomBC-2, $KomB^mC$ and $KomBC^m$ were heated to 90 °C for 10 min and then centrifuged at 13000 g at 4 °C for 15 min to remove precipitated protein. The supernatant was filtered through 0.22 µm membrane filters for LC-MS/MS analysis. For the latter, the reaction of KomA and *E. coli* NDK with dADP and ATP was performed as described in the **εNAD-based NADase assay**, except that the 50-µl reaction mixtures contained 1 mM dADP, 0 or 20 µM KomA, 2 mM ATP and 0 or 4 µM NDK. Then the mixtures were heated to 90 °C for 5 min to stop the reaction, and supplemented with 150 µl MilliQ $H_2O$. The mixtures were then centrifuged at 13000 g for 10 min to remove precipitated protein and the supernatants was passed through 0.2 µm Filters for LC-MS/MS analysis. The reaction of R1DNK with dIMP and ATP was performed as described in the **εNAD-based NADase assay**, except that the 100-µl reaction mixtures contained 1 mM dIMP, 5 µM R1DNK, 3 mM ATP. Then the mixtures were heated to 90 °C for 5 min to stop the reaction. The mixtures were then centrifuged at 13000 g for 10 min to remove precipitated protein and the supernatants was passed through 0.2 µm Filters for LC-MS/MS analysis.

The LC–MS/MS analysis was performed using an UltiMate 3000 UHPLC System coupled to the Q Exactive Plus Orbitrap high-resolution mass spectrometry (Thermo Fisher Scientific). The UHPLC was performed using an InfinityLab PoroShell 120 HILIC-Z PEEK lined column (Agilent Technologies, 2.1 × 100 mm, particle size 2.7 µm). The column was warmed to 30 °C and the flow rate was 0.3 ml/min. Three µl of each sample was loaded onto the column. Gradient elution was performed with solvent A (10 mM ammonium acetate at pH 9 in ultra-pure water) and solvent B (10 mM ammonium acetate at pH 9 in 90% acetonitrile and 10% ultra-pure water). The gradient elution proceeded as follows: 0 to 12 min, 90% to 50% B; 12 to 13 min, 50% B; 13 to 13.1 min, 50% to 90% B; 13.1 to 26 min, 90% B.

MS/MS analysis was performed in negative ionization mode from 100 to 1200 m/z at a mass resolution of 70,000. Synthetic dITP (Thermo Fisher Scientific, R1191) was also run to identify the dITP standard peaks. The reaction product of KomA and dADP was used as dIDP standard. MS/MS spectra collection was performed at a resolution of 17,500. Data analysis was performed using Xcalibur Qual Browser (Thermo Fisher Scientific).

**Quantification of nucleotides by UHPLC-MS/MS**

To analyze the nucleotides in cells infected by phages, overnight cultures of *E. coli* MG1655 were diluted 1:100 in 200 ml MMB medium and grown to an $OD_{600}$ of 0.3 at 37 °C. 45 ml of cultures were sampled (Unifected), and then the rest cultures were infected with WT phage or mutants at a final MOI of 3. Subsequently, 45 ml of infected cultures were taken at indicated times (5 min, 10 min, and 15 min for T5 or 8 min, 16 min, and 24 min for Rao1) and centrifuged (8000 g, 4 °C) for 5 min to remove supernatants.

To analyze the nucleotides in cells expressing Kongming and R1DNK, the mutated system (KomABC$^m$) was expressed from its native promoter, while R1DNK was expressed from an arabinose promoter. The cells were collected after the expression of R1DNK was induced for 1 hour.

The pellets were resuspended in 700 µl of cold extraction solvent (acetonitrile, methanol, water, formic acid = 2:2:1:0.02, v/v/v/v) and lysed by ultrasonication for 1 min, followed by centrifugation at 13000 g at 4 ℃ for 15 min to remove the debris. The supernatants were dried by lyophilization, then resuspended in 100 µl water and analysed by UHPLC-MS/MS. For analyses of the nucleotides in cells expressing R1DNK, the method was similar to that described above, except that MMB medium was supplemented with 0.2% arabinose and corresponding antibiotic and samples were taken at 60 min post-induction.

Quantification of nucleotides were performed using an UltiMate 3000 UHPLC System coupled to a TSQ Plus triple quadrupole mass spectrometer (Thermo Fisher Scientific). The UHPLC was performed as described in the **Liquid chromatography-tandem mass spectrometry.** The analyses were performed in positive ion modes (dATP, dADP, dAMP) or negative ionization mode (dIMP, dIDP, dITP). Nucleotides were detected and quantified using multiple reaction monitoring (MRM).

A standard curve of dITP was constructed within the concentration range from 5 µM to 60 µM for quantification of the released dITP from the denatured protein samples. Metabolites (dATP, dADP, dAMP, dIMP, dIDP, and dITP) were quantified using product ion area. Data analysis was performed using Xcalibur Qual Browser (Thermo Fisher Scientific).

**Identification of DNK/Dmp homologues**

Phage DNK and Dmp homologues were identified in the INPHARED database (release 1Mar2024)(*9*). The protein set used to construct HMM models for DNK and Dmp were identified using HMMER v3.4's jackhmmer (*10*) with an E-value of 1e-3, employing T5 DNK (YP_006871.1) and Dmp (YP_006829.1) sequences as queries against our phage collection database. HMM profiles were generated using HMMbuild after removal of redundant sequences by CD-HIT v4.8.1 (*11*) with sequence identity threshold 100% and length difference cutoff 100%. DNK and Dmp homologues were identified using the generated HMM profiles as query against INPAHARED database of RefSeq complete phage genomes with an E-value threshold of 1e-3.

**Phylogenetic tree construction**

To construct phage phylogenetic tree, the phages used in this study were clustered using Genome-BLAST Distance Phylogeny (GBDP) method with the D0 formula for nucleotides in VICTOR (*12*). Taxonomy information was retrieved from NCBI database or BASEL phage collection (*1*).

KomB domain-containing proteins were retrieved using HMMER v3.4's hmmsearch with the Pfam HMM profile PF01725 as the query against the RefSeq non-redundant protein database. Proteins with sequences longer than 100 amino acids were clustered using MMseqs2 easy-cluster (*13*). Representative sequences were aligned using hmmalign with the '-trim' flag, and a phylogenetic tree was constructed using FastTree v2.1.11 (*14*) with gamma-distributed site rates and the WAG evolutionary model.

For other phylogenetic trees, sequences were first aligned with MAFFT v7.525 FFT-NS-1000 (*15*) and trimmed using trimAl v1.5.0 (*16*) to remove any column in the alignment that has more than 30% gaps. Phylogenetic tree construction was performed using FastTree with gamma-distributed site rates and the WAG evolutionary model. Tree visualized using the Python package etetoolkit (*17*) or ITOL (*18*).

**Sequence analysis and structure prediction**

Sequence alignment of KomA (VFT02523), KomB (VFT02524), and KomC (VFT02525) with their respective homologs was performed with Jalview (fig. S2)(*19*). Structures of the three proteins were predicted using AlphaFold3 (*20*). The predicted structures were aligned with the experimentally determined structures of their respective homologs using ChimeraX v1.8 (*21*).

To perform sequence logo analysis, all HAM1 protein sequences retrieved as described above were aligned using FAMSA v2.2.2 (*22*) with UPGMA tree mode, and the aligned fasta sequences were extracted separately for indicated clades. The sequence logo for each part was generated using Weblogo v3.7.12 (*23*) with chemistry color-scheme.

**Genomic context analysis of *ham1* genes**

To determine whether a *ham1* gene is located within a defense island, known defense systems were identified using DefenseFinder v1.2.2 (*24*) and PADLOC v2.0.0 (*25*) within 20,000 bp upstream and downstream of the *ham1* gene. Then, the percentage of *ham*1 genes with defense systems located within ±20,000 bp was calculated for each specific HAM1 clade.

To analyze whether HAM1 is encoded near or in prophages and plasmids, the genetic elements were detected using the geNomad v1.8.0 (*26*) end-to-end pipeline within 75,000 bp upstream and downstream of the *ham1* gene. If prophages or plasmids were detected, the *ham1* genes were counted as being within or near mobile genetic elements.

To analyze the domains of the proteins encoded in ham1 neighborhood, hmmsearch is used to identify and annotate protein domains by comparing protein sequences against a library of HMM profile.

**Spearman Analysis**

The 16S rRNA phylogenetic tree was constructed for all organisms encoding defensive HAM1s, in which the 16S SSU rRNA, HAM1, and associated Sir2 genes were found. The 16S rRNA genes were identified via BLASTN v2.15.0+ searches (*27*), using the 16S rRNA sequence of *Escherichia coli* K-12 ER3413 as the query against bacterial genomes, with a word size of 8 and dust filtering disabled. Hits with at least 80% coverage and 70% identity were selected and aligned using MAFFT v7.525, and a phylogenetic tree was constructed using FastTree v2.1.11 with gamma-distributed site rates and the GTR evolutionary model. HAM1, and Sir2 protein sequences were aligned using MAFFT FFT-NS-1000, and approximate maximum-likelihood trees were generated using FastTree with gamma-distributed site rates and the WAG evolutionary model.

**Statistics Analysis**

For all statistics analysis, the assay were performed in 3 independent replicates as indicated in the figure legends, and the unpaired t-test was used to calculate p values.

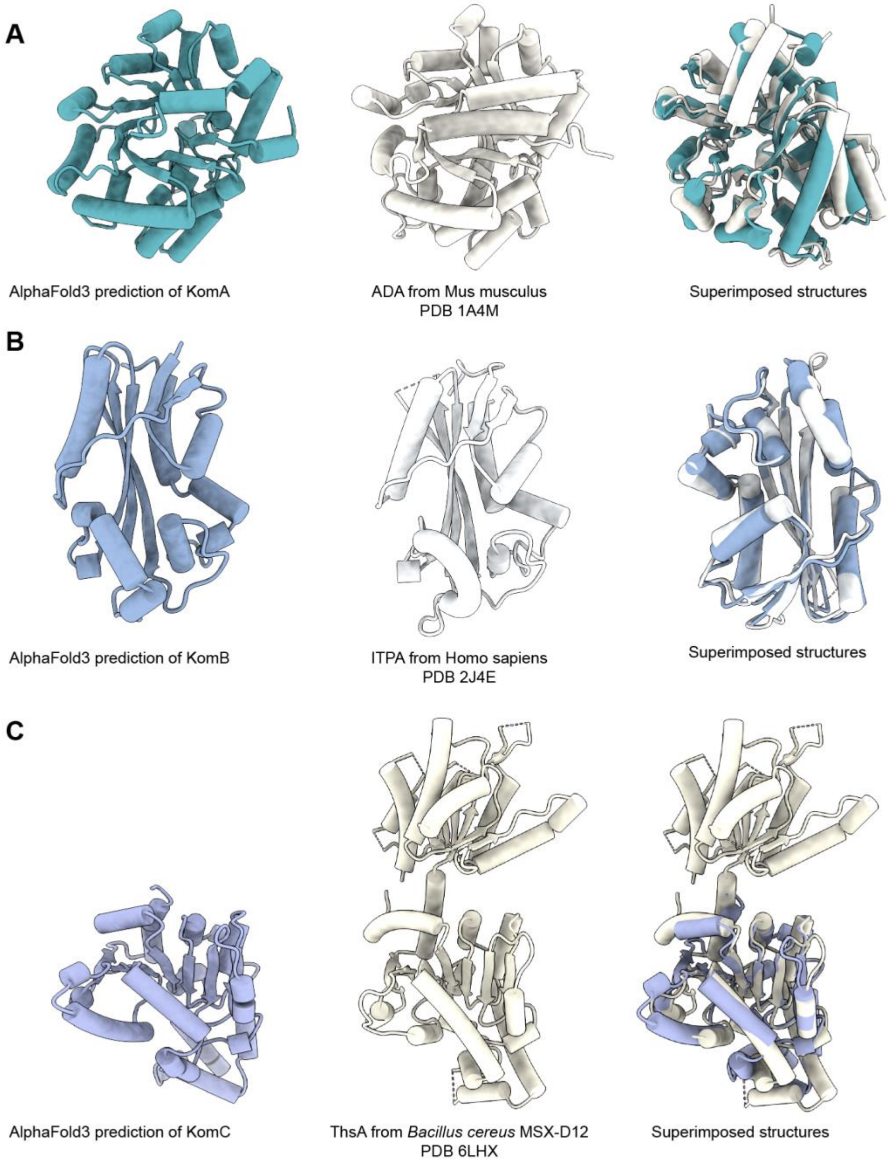


Fig. S1 Structural homology of the proteins encoded by the three-gene operon to characterized enzymes. The structures of the KomA (A), KomB (B) and KomC (C) proteins from *Escherichia coli* NCTC13216 are predicted by AlphaFold3 (left panels). The experimentally determined structures of *Mus musculus* adenosine deaminase (ADA, PDB: 1A4M), *Homo sapiens* ITPA (PDB: 2J4E) and *Bacillus cereus* MSX-D12 ThsA (PDB: 6IHX) are shown in middle panels, and the superimposed structures are shown in the right.

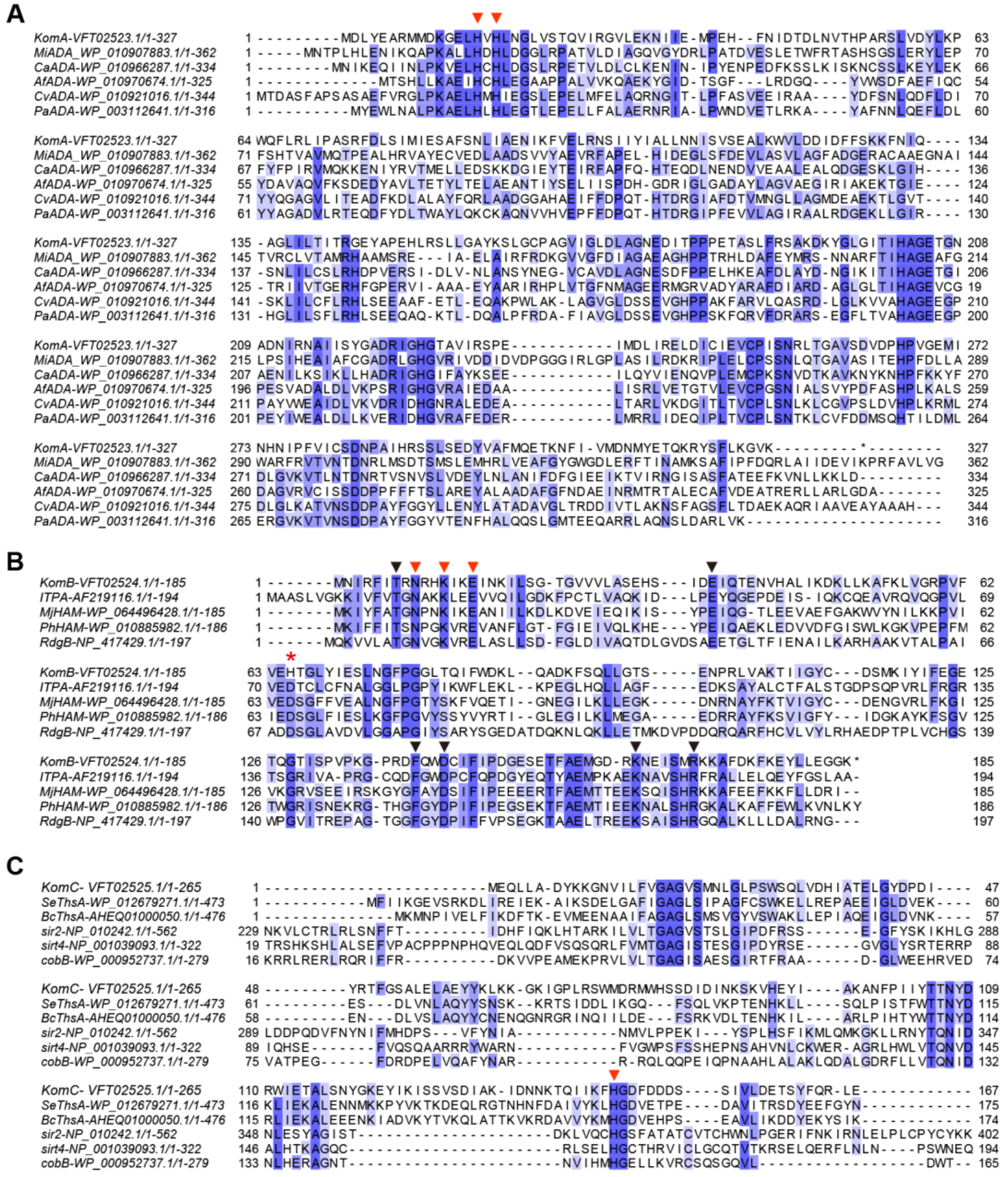


Fig. S2 Sequence alignment of the three proteins of the Kongming and their representative homologues respectively. (A) KomA; (B) KomB; (C) KomC. In (B), the red star indicates that the conserved catalytic aspartic acid in housekeeping HAM1 family enzymes is replaced by histidine in the HAM1 of KomABC (His$^{65}$), while the arrows, either red or black, indicate the conserved residues implicated in non-canonical purine NTP substrate binding. In (A-C), the residues marked with red arrows were subjected to mutagenesis: KomA, alanine substitution of His$^{15}$ and His$^{17}$ (H15A-H17A); KomB, alanine substitution of Asn$^{9}$, Lys$^{12}$ and Glu$^{15}$ (N9A-K12A-E15A); KomC, alanine substitution of His$^{146}$ (H146A).

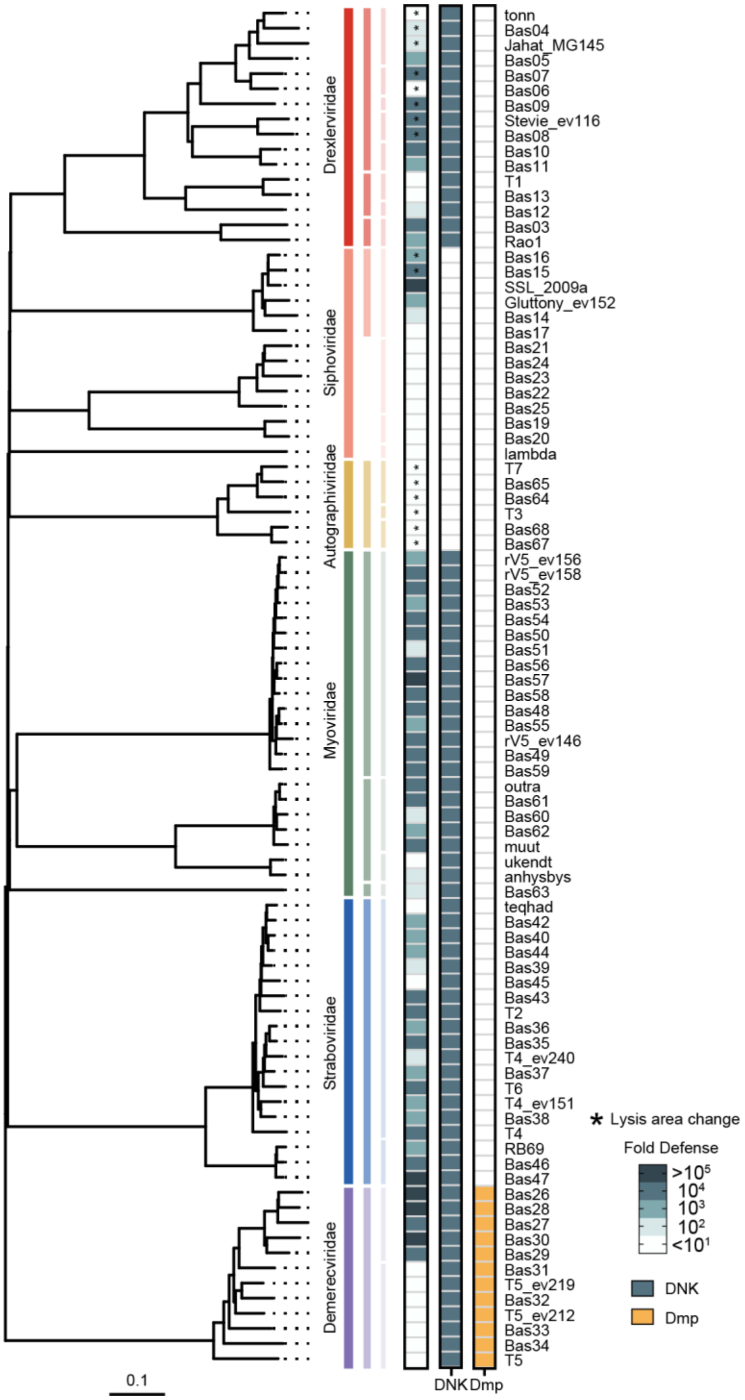


Fig. S3 Fold protection of Kongming against the phage collection in our lab. The distributions of DNK and Dmp homologues are indicated.

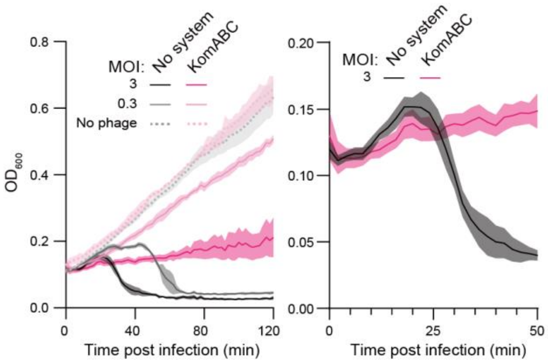


Fig. S4 Kongming induces growth retardation earlier than the control strain post Rao1 infection at MOI 3. Left panel: a copy of Fig. 1D; right panel: magnified view of Fig. 1D.

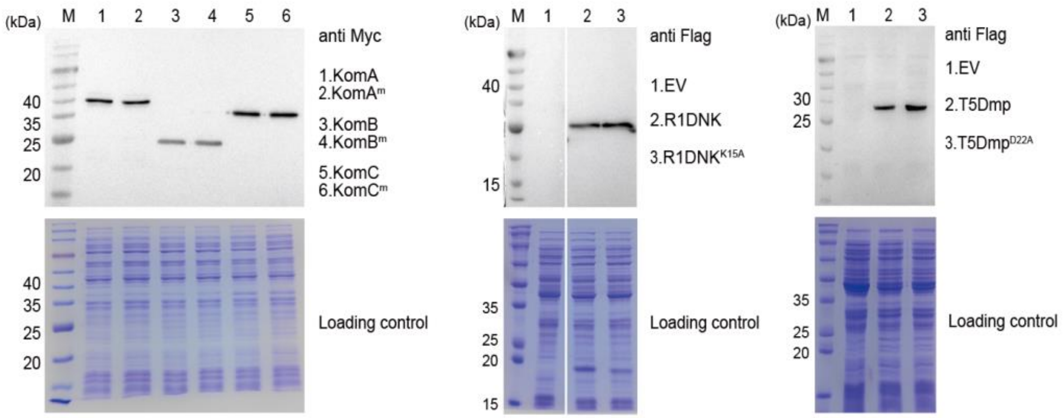


Fig. S5 The mutations of the proteins analyzed in this study do not affect their expression level. Cell lysates were prepared from the cultures expressing the indicated proteins and their mutants, and analyzed by western blot with the indicated antibody. The cell lysates were also analyzed by SDS-PAGE as loading control. EV, cultures carrying an empty vector.

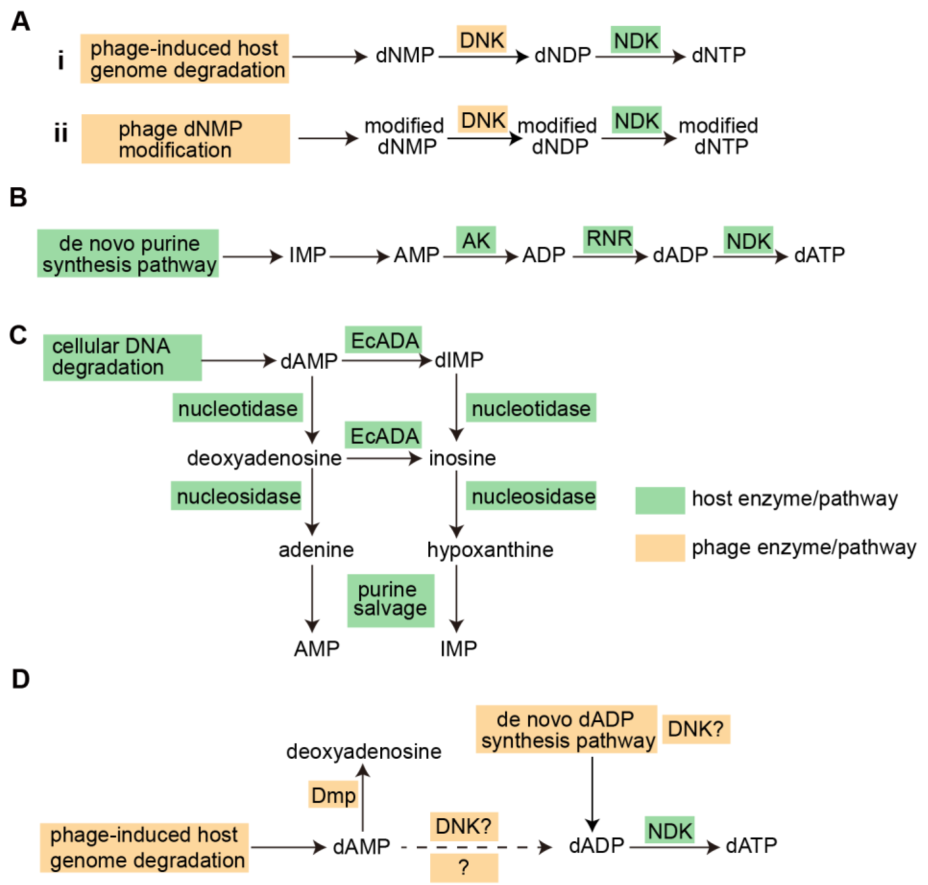


Fig. S6 Schematic of host and phage nucleotide metabolsim.

(A) Proposed functions of phage DNKs. (i) Phage DNKs phosphorylate dNMP that are generated by host genome degradation to dNDK, which can be used to increase the dNTP pool to facilitate phage genome replication. (ii) Phage DNKs phosphorylate motified dNMP, which is converted to motified dNTP and used for synthesis of motified genome. The modification renders phage genome resistant to many nuclease-based defense systems.

(B) Host dNTP synthesis pathway with dATP as an example. The de novo purine synthesis pathway produces IMP, which is then converted to AMP, followed by generation of ADP by adenylate kinase (AK). Reduction of ADP by ribonucleotide reductase (RNR) produces dADP, which is converted to dATP by nucleoside diphosphate kinase (NDK).

(C) Recycle of purine from dAMP. dAMP may be produced by DNA degradation that occurs during DNA replication and repair. Host housekeeping adenosine deaminase (EcADA) converts dAMP to dIMP, and both dAMP and dIMP can be degraded into bases that are then recycled in the purine salvage pathway.

(D) Nucleotide metabolism of T5. T5 infection induces extensive degradation of host genome and the generated dNMP is rapidly cleared by DMP. Upon deletion of *dmp*, the dAMP can be converted to dADP by DNK and/or an alternative unknown mechanism (dashed line). T5 encodes its own de novo dNTP synthesis enzymes, including RNR to produce dADP. Whether DNK plays a role in the T5 de novo dNTP synthesis remains unclear.

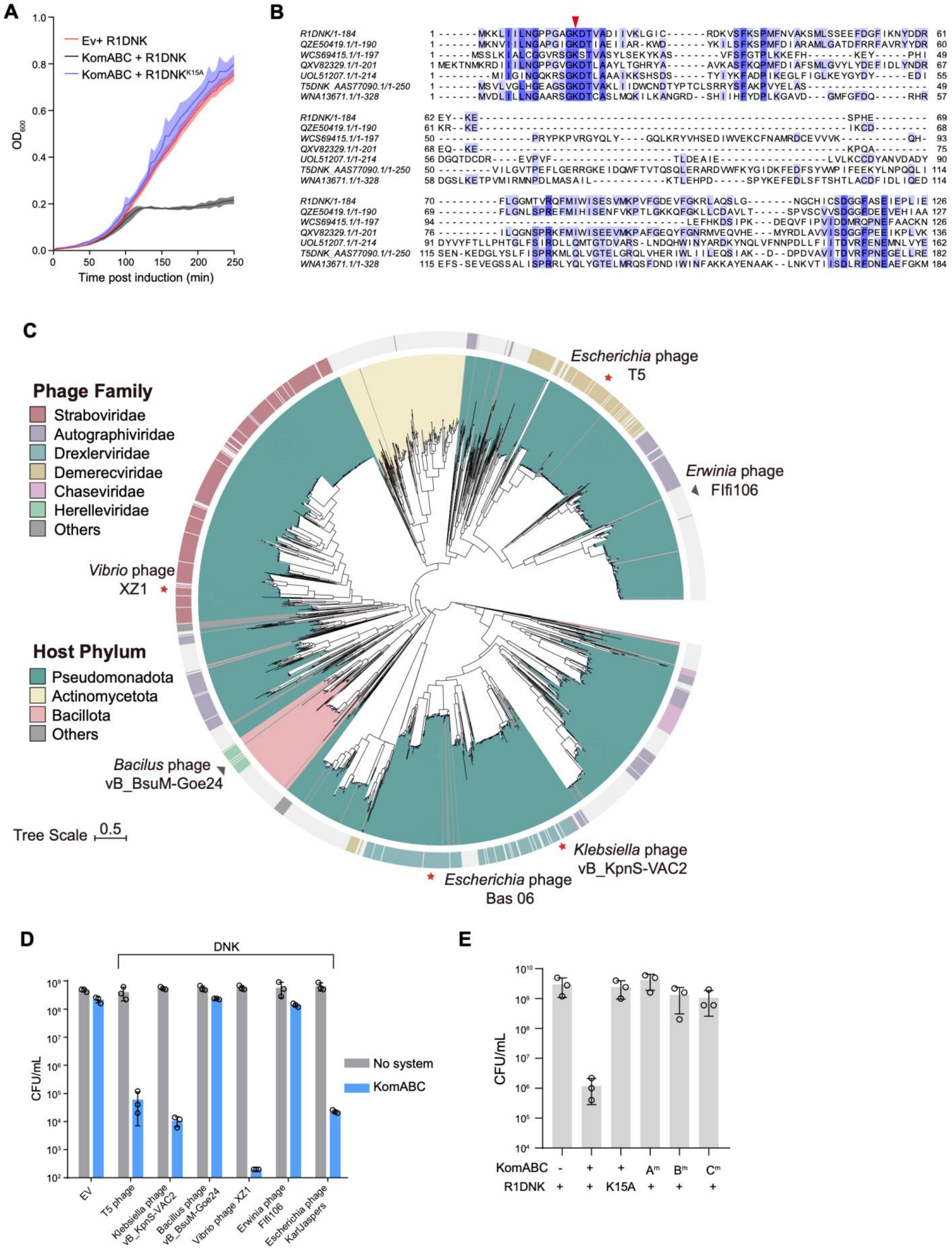


Fig. S7 Phage DNKs trigger Kongming to mediate toxicity.

(A) Co-expression of R1DNK and the system (KomABC) suppresses culture growth. The optical density at 600 nm ($OD_{600}$) was measured after the expression of R1DNK or R1DNK$^{D22A}$ was induced by arabinose in cells expressing KomABC or an empty vector. Data represent the mean ± standard deviation of n = 3 biological replicates.

(B) Sequence alignment of R1DNK and its representative homologues. The catalytic site of the DNKs is indicated by an arrow.

(C) Enlarged view of phage DNK phylogenetic tree.

(D) Six DNK homologues were tested for their capability to trigger Kongming. Colony formation units (CFUs) were measured for cells co-expressing KomABC and DNKs from the indicated phages. Data represent the mean ± standard deviation of n = 3 biological replicates, with individual data points overlaid.

(E) The cultures used for the analysis of $NAD^+$ depletion (Fig. 2A) were grown to 2 hours post induction of R1DNK expression, and then, colony formation units (CFUs) were measured. Data represent the mean ± standard deviation of n = 3 biological replicates, with individual data points overlaid.

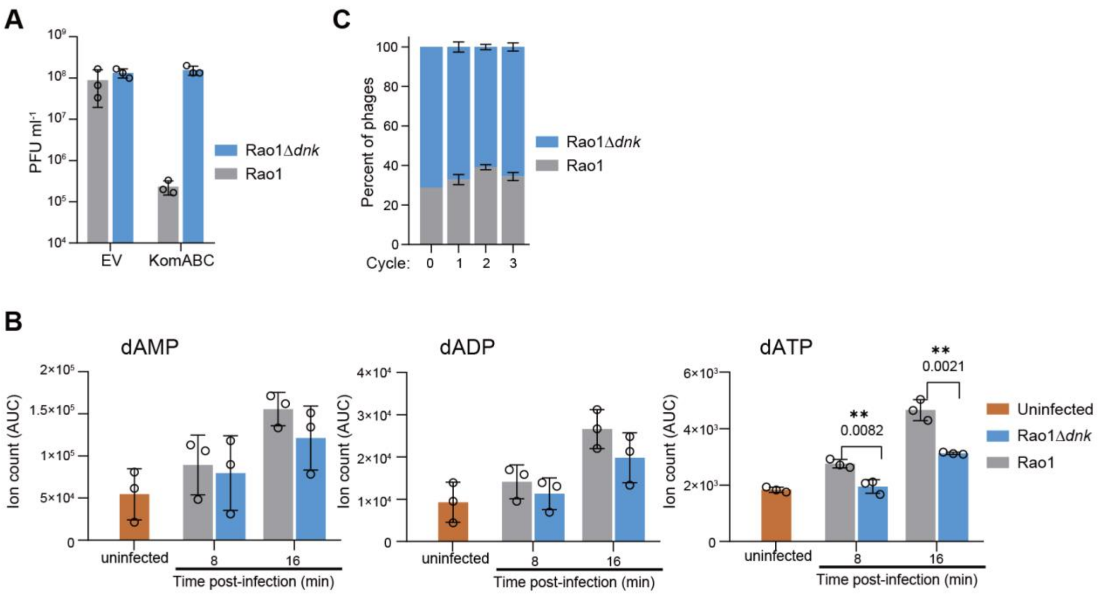


Fig. S8 Characterization of *dnk* deletion Rao1 mutant.

(A) Plaque formation units of wild type Rao1 and Rao1$^{\Delta dnk}$ on cells carrying empty vector (EV) or Kongming (KomABC).

(B) Ion count (area under curve) of dAMP, dADP and dATP in lysates extracted from cells infected with Rao1 and Rao1$^{\Delta dnk}$, or uninfected cells. The cell lysates were prepared at 8 and 16 min post infect at MOI of 3, and the nucleotides were analyzed by LC-MS/MS. Data represent the mean ± standard deviation of n = 3 biological replicates, with individual data points overlaid.

(C) Phage competition fitness assay between wild type Rao1 and Rao1$^{\Delta dnk}$. The y axis represents the fraction of each phage of both phages based on sequenced DNA reads. The x axis represents the infection cycle, with 0 representing the mixture prior to first infection cycle.

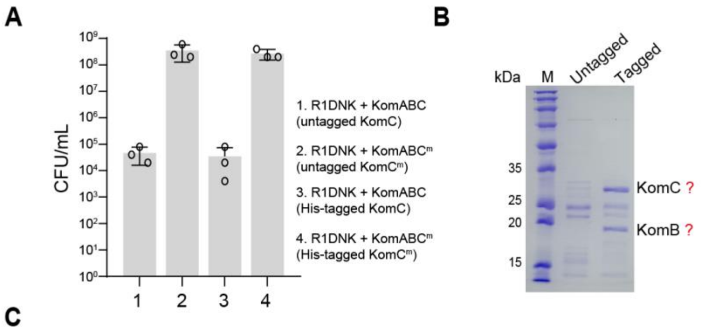


C

| Accessions | Description | Gene | MW [kDa] | Abundances (Normalized)_Gel Slice 1 |
|---|---|---|---|---|
| VFT02524.1 | deoxyribonucleotide triphosphatepyrophosphatase | NCTC13216_03967 KomB | 20.9 | 1818287805 |
| WP_000719886.1 | Sel1 family TPR-like repeat protein YjcO | yjcO | 25.1 | 275367656 |
| WP_000673804.1 | peptidoglycan glycosyltransferase_ peptidoglycan DD-transpeptidase MrcA | mrcA | 93.5 | 196774656 |
| WP_001269673.1 | LPS assembly lipoprotein LptE | lptE | 21.3 | 47462512 |
| WP_000861334.1 | peptidylprolyl isomerase | slyD | 20.8 | 3321716 |
| VFT02525.1 | Sir2 family NAD-dependent protein deacetylase | NCTC13216_03968 KomC | 30.6 | 2993202.25 |

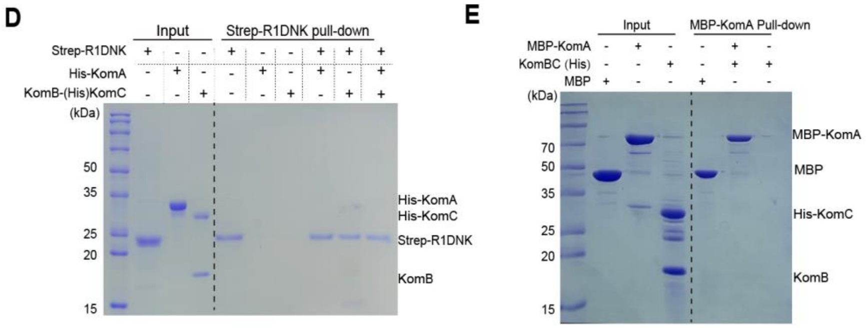


Fig. S9 Analysis of the physical interaction of R1DNK and the three proteins of Kongming.

(A) His-tag within KomC does not affect the function of Kongming. Colony forming units (CFUs) were measured for cells co-expressing R1DNK and KomABC or its variants carrying KomC mutation (KomC$^m$, H146A) or His-tag. Data represent the mean ± standard deviation of n = 3 biological replicates, with individual data points overlaid.

(B) SDS-PAGE analysis of the co-purification with untagged KomC$^m$ and His-tagged KomC$^m$ from cell expressing R1DNK and KomABC$^m$. The bands putatively corresponding to His-tagged KomC and KomB are indicated respectively.

(C) Mass spectrometry analysis of the co-purified band with His-tagged KomC from cells expressing KomC and KomB (Fig. 2C). Top six proteins with the highest normalized abundance are shown.

(D-E) SDS-PAGE analysis of the pull-down assay using Strep-R1DNK to capture His-KomA and/or KomB-(His)KomC complex (D), or using MBP-KomA to capture KomB-(His)KomC complex (E). On the left side of the dashed line are the proteins used for the pull-down assay; on the right are the proteins bound to the Strep-Tactin resin (for Strep-R1DNK) or Dextrin Beads (for MBP-KomA).

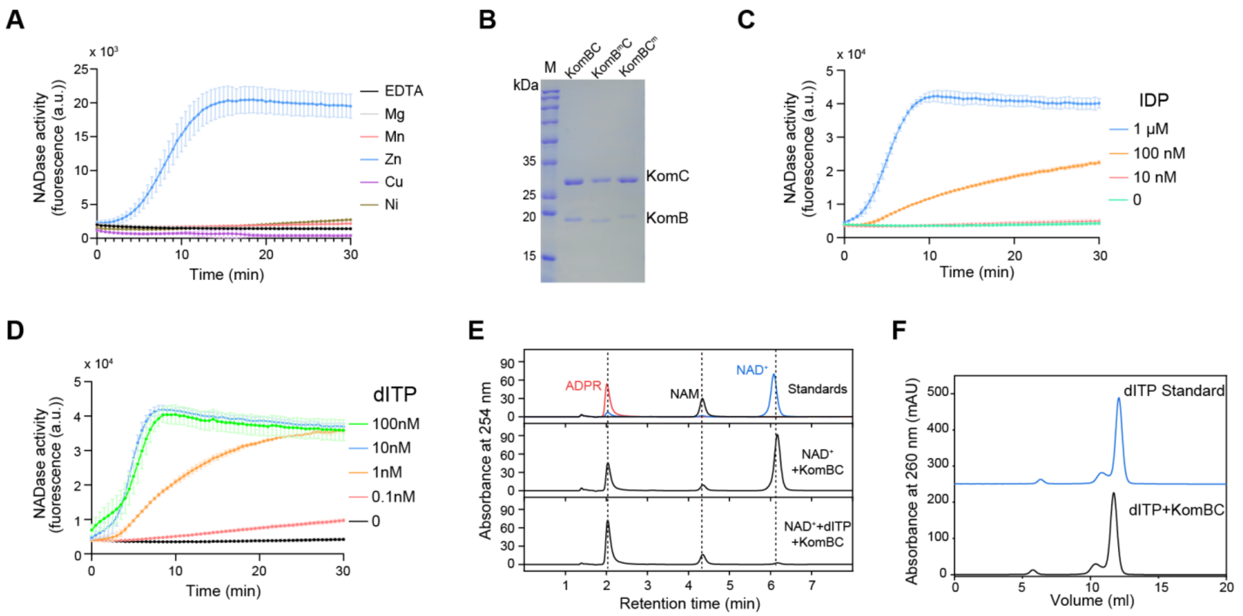


Fig. S10 Characterization of the activities of KomB-KomC complex (KomBC).

(A) NADase activity of KomBC in the presence of EDTA or indicated divalent metal ions.

(B) SDS-PAGE analysis of purified KomBC and the mutated complexes. $B^{m}$, $KomB^{N9A\text{-}K12A\text{-}E15A}$; $C^{m}$, $KomC^{H146A}$.

(C) NADase activity of KomBC in the presence of a gradient of IDP.

(D) NADase activity of KomBC in the presence of a gradient of dITP.

(E) High-performance liquid chromatography (HPLC) analysis of the $NAD^{+}$ degradation products by KomBC in the presence of dITP. The experiments were performed simultaneously with those in Fig. 2E, and the same standards are shown.

(F) Ion-exchange chromatography analysis of the pyrophosphatase activity of KomBC. dITP was incubated with KomBC and then the reaction mixture was analyzed by ion-exchange chromatography with dITP as standard.

All experiments were performed three independent times, and data represent the mean ± standard deviation of n = 3 biological replicates for the NADase assay.

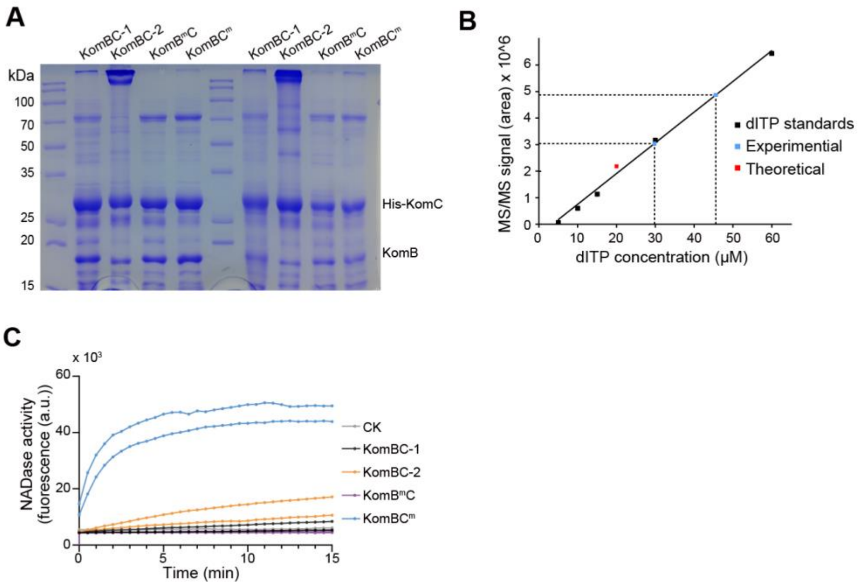


Fig. S11 Recycle of signaling molecule from purified KomBC$^{m}$ complex.

(A) SDS-PAGE analysis of the protein samples from Fig. 3C.

(B) LC-MS/MS calibration curve of dITP standard (marked in black) and the observed amount of dITP (marked in blue) in the supernatant from the denatured KomBC$^{m}$ complex from Fig. 3C. The red dot (Theoretical) indicates the measurement of 20 μM dITP standard using the curve

(C) NADase activity of 50 nM KomBC in the presence of the released molecules from the denatured protein samples from Fig. 3C. The experiments were performed in two independent replicates. CK: without any released molecules.

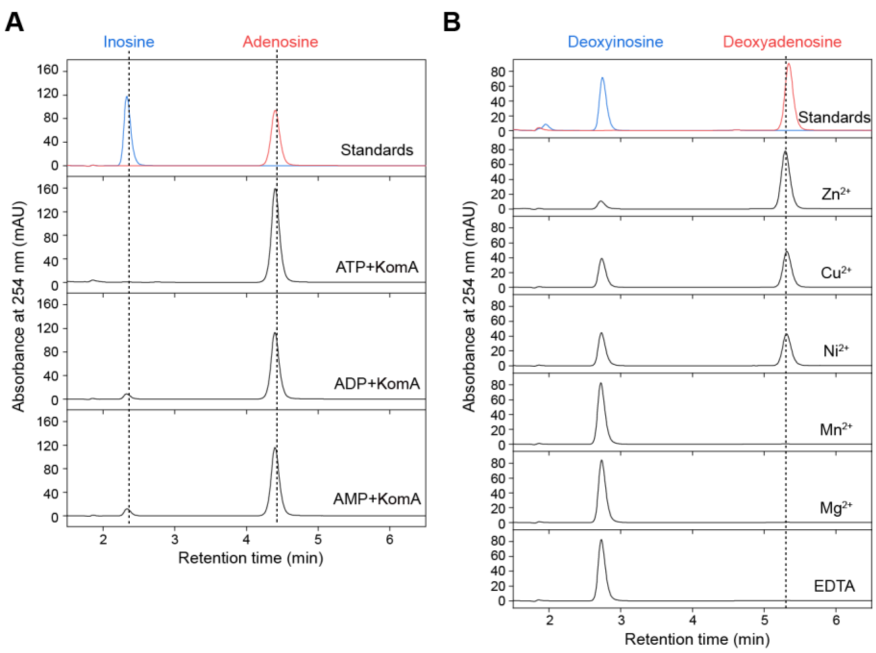


Fig. S12 Characterization of the activity of KomA.

(A) HPLC analysis of the adenosine deaminase activity of KomA with ATP, ADP or AMP as substrates. Deoxyadenosine and deoxyinosine standards are shown at the bottom.

(B) Effects of EDTA and divalent metal ions on the adenosine deaminase activity of KomA with dAMP as substrate. All experiments were performed three independent times.

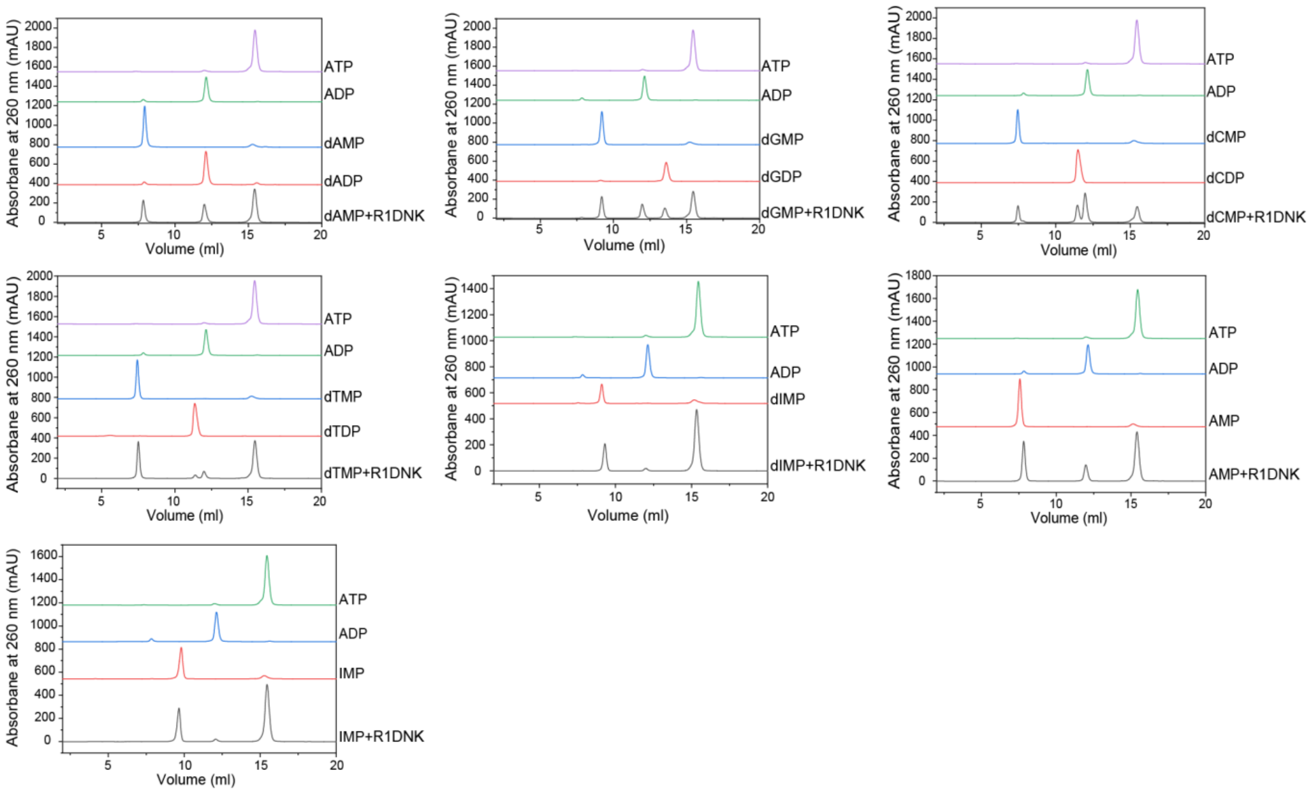


Fig. S13 Nucleotide kinase assays of R1DNK with all standards. The tested substrates include dAMP, dGMP, dCMP, dTMP, dIMP, AMP and IMP. All experiments were performed three independent times.

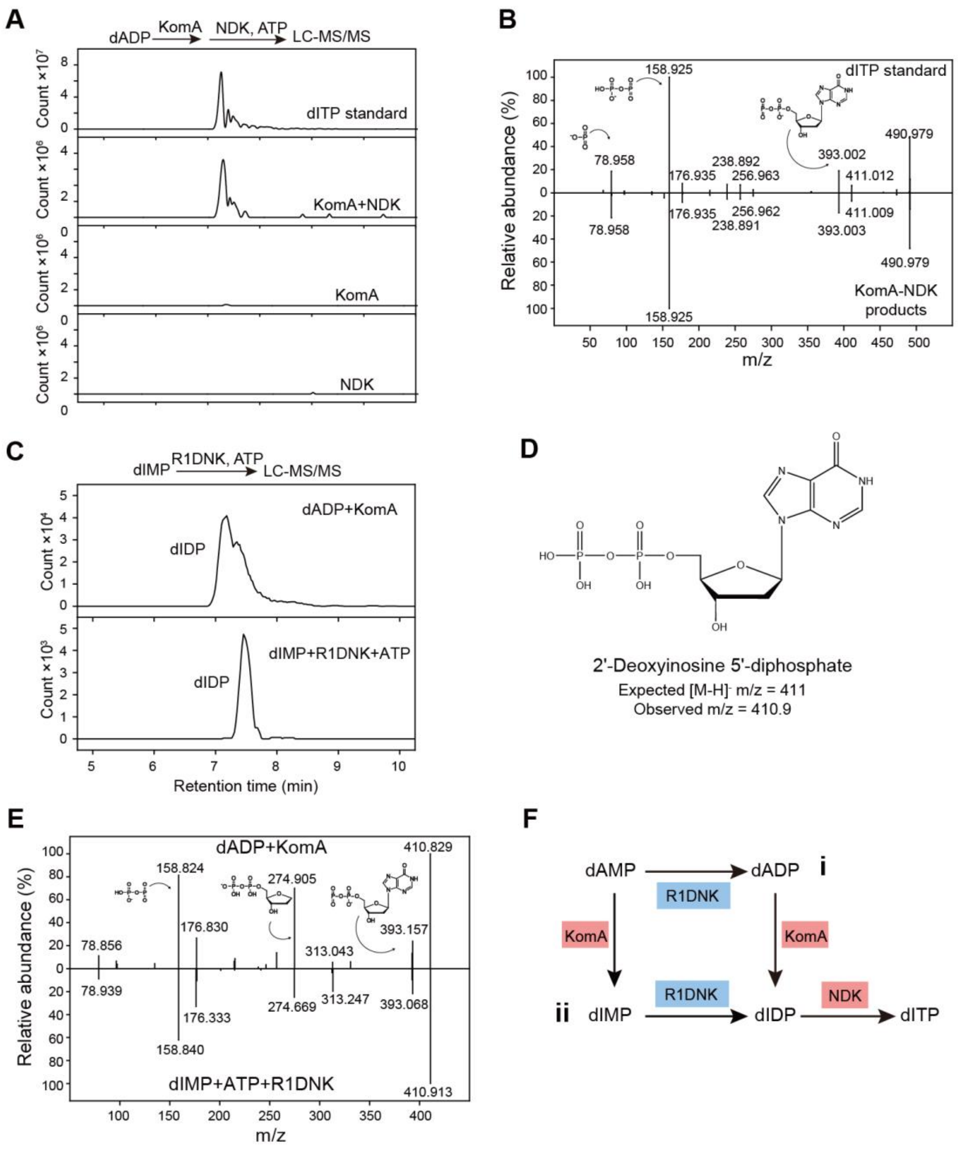


Fig. S14 Analysis of the reactions with KomA, NDK and R1DNK using LC-MS/MS.

(A) LC-MS analysis of dITP standard and the reaction product of KomA and/or NDK with dADP and ATP. Extracted mass chromatograms of ions with an m/z value of 490.97 are presented.

(B) MS/MS analysis of dITP standard and the reaction product of KomA and NDK with dADP and ATP.

(C) LC-MS analysis of the reaction product of KomA and dADP (as standard) and the reaction product of R1DNK, dAMP and ATP. Extracted mass chromatograms of ions with an m/z value of 410.9 are presented.

(D) Chemical structure of dIDP and its m/z value.

(E) MS/MS analysis of dIDP standard (KomA and dADP) and the reaction product of R1DNK, dAMP and ATP.

(F) Proposed two dITP synthesis pathways.

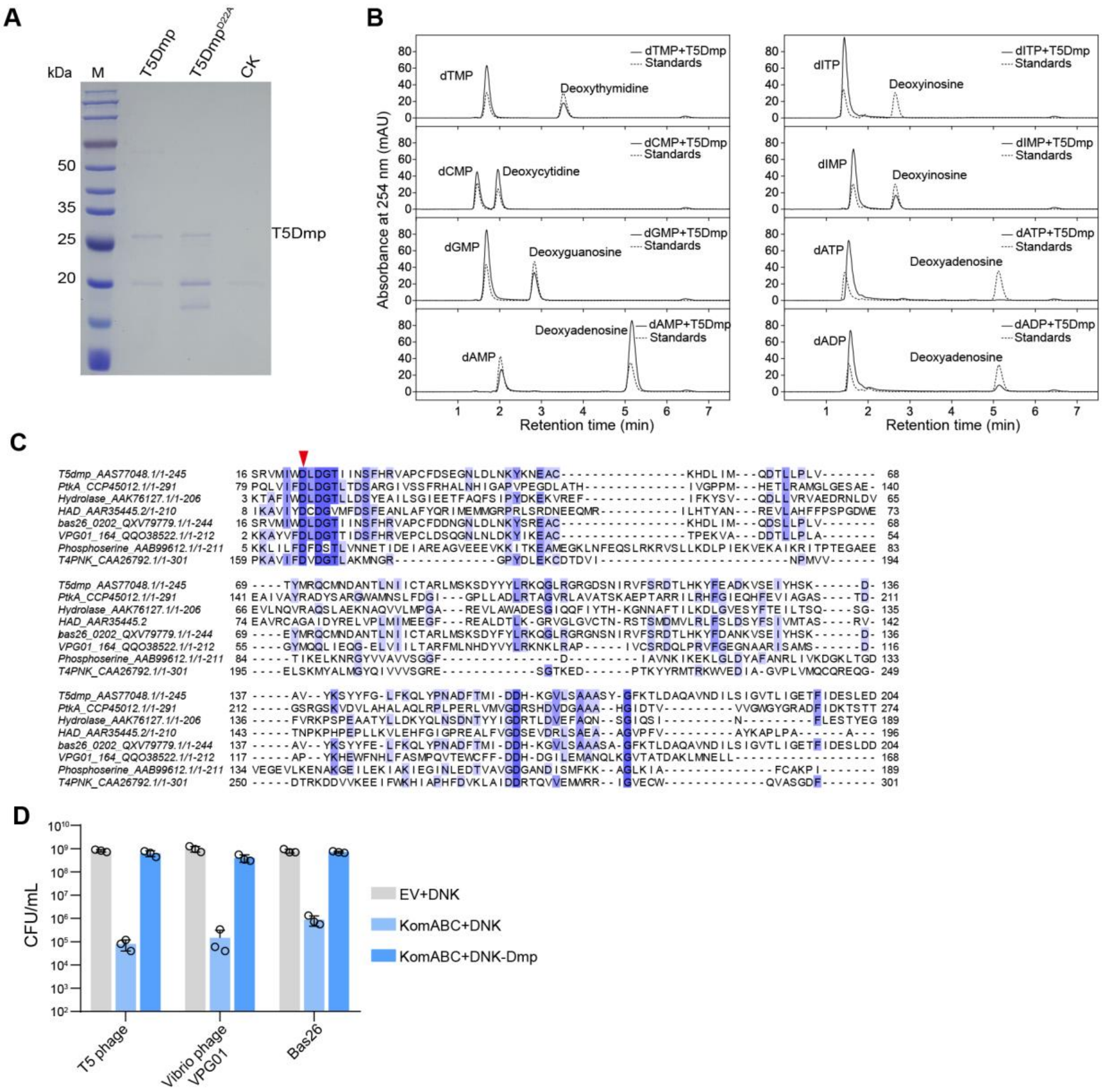


Fig. S15 Characterization of T5Dmp and its homologues.

(A) SDS-PAGE analysis of purified T5Dmp and a D22A mutant. CK, control purification from cells containing an empty vector.

(B) The phosphatase activity of T5Dmp with dTMP, dCMP, dGMP, dAMP, dITP, dTMP, dIMP, dATP or dADP as substrate. All experiments were performed three independent times.

(C) Sequence alignment of T5Dmp and other representative phosphatases. The catalytic site of the phosphatases is indicated by an arrow.

(D) Two T5Dmp homologues, as well as T5Dmp, were tested for their capability to inhibit KomABC. Colony formation units (CFU) were measured for cells co-expressing KomABC and DNKs, or co-expressing KomABC, DNKs and Dmps from the indicated phages. EV, an empty vector control for KomABC. Data represent the mean ± standard deviation of n = 3 biological replicates, with individual data points overlaid.

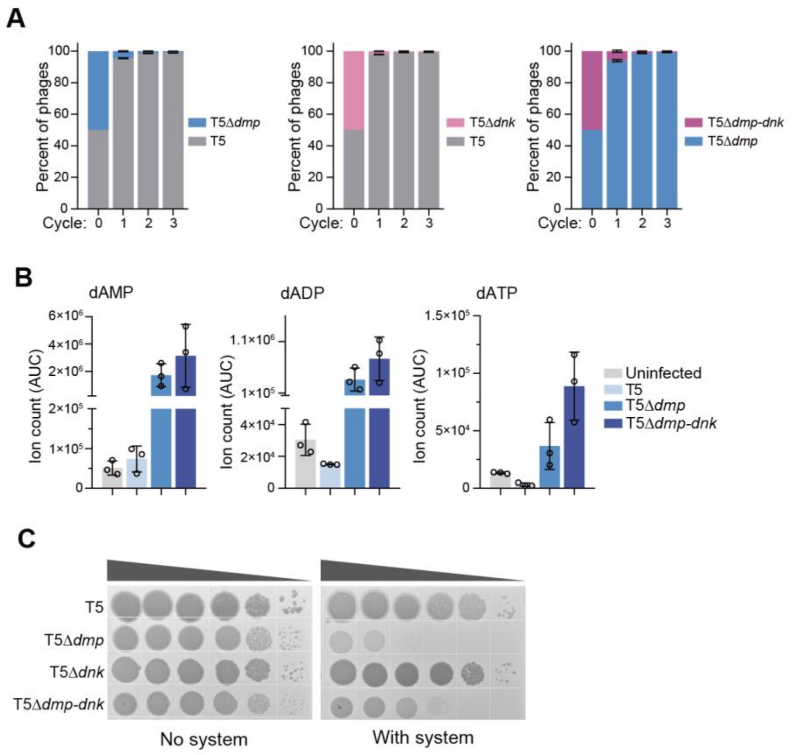


Fig. S16 Characterization of *dmp* and *dnk* deletion T5 mutants.

(A) Phage competition fitness assay between wild type T5 and T5$^{\Delta dmp}$ (left panel), T5 and T5$^{\Delta dnk}$ (middle panel), T5$^{\Delta dmp}$ and T5$^{\Delta dmp\text{-}dnk}$ (right panel). The y axis represents the fraction of each phage of both phages based on sequenced DNA reads. The x axis represents the infection cycle, with 0 representing the mixture prior to first infection cycle.

(B) Ion count (area under curve) of dAMP, dADP and dATP in lysates extracted from cells infected with T5, T5$^{\Delta dmp}$ and T5$^{\Delta dmp\text{-}dnk}$, or uninfected cells. The cell lysates were prepared at 10 min post infect at MOI of 3, and the nucleotides were analyzed by LC-MS/MS. Data represent the mean ± standard deviation of n = 3 biological replicates, with individual data points overlaid.

(C) Plaque formation units of wild type T5, T5$^{\Delta dmp}$, T5$^{\Delta dnk}$ and T5$^{\Delta dmp\text{-}dnk}$ on cells expressing Kongming or not.

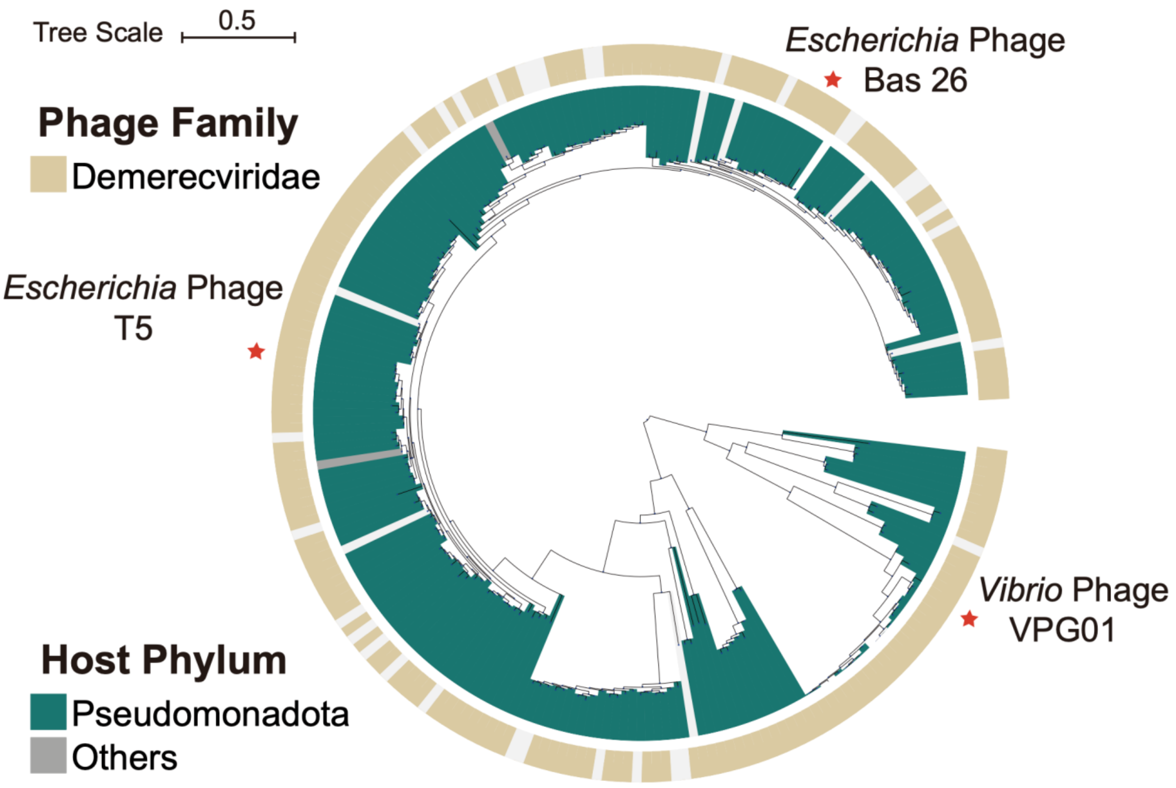


Fig. S17 Phylogenetic analysis of phage-encoded Dmps retrieved from the RefSeq complete phage genomes database (INPHARED). The phages encoding the Dmps experimentally validated to harbor anti-Kongming function are indicated by red stars.

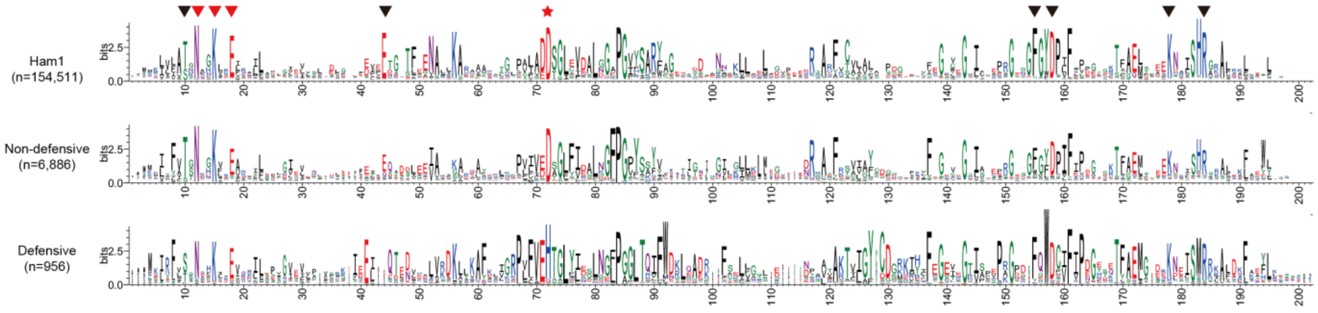


Fig. S18 Sequence logo representation of all, non-defensive and defensive HAM1s respectively. The red star indicates the substitution of the conserved catalytic aspartic acid in housekeeping HAM1s by histidine in defensive HAM1s, while the arrows indicate the conserved residues implicated in non-canonical purine NTP substrate binding

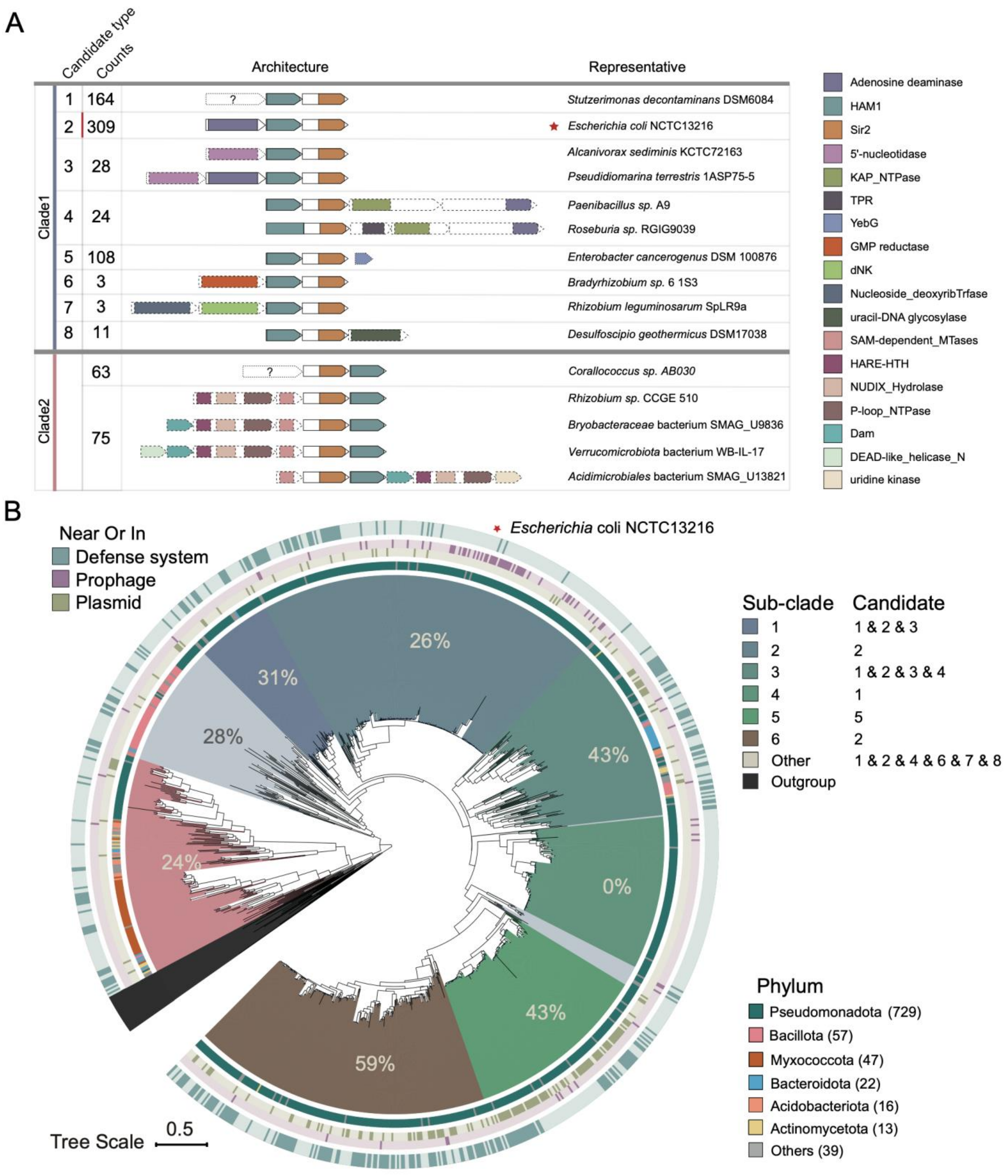


Fig. S19

(A) Domain organization of predicted systems containing defensive HAM1. The system candidates are classified by the associated proteins and their operon architectures. Candidates in broken contigs are not considered.

(B) Phylogenetic tree of defensive HAM1s showing subclades of clade 1. For each subclade, the percentage of HAM1s that are encoded near known defense systems is indicated. Subclades and candidate types correlation are shown on the right. From outer to inner circles: whether each HAM1 is encoded near known defense systems, near or in prophages, near or in plasmids, and the bacterial phylum to which the HAM1 belongs. Phylum comprising less than 1% are classified as "others". Sequences without taxon info are not considered.

**Table S1 Proteins analyzed in the study. The names, accession numbers, source strains, and sequences are listed.**

**Table S2 Plasmids used in the study.**

**Table S3 Primers used in the study**

**Table S4 Phages used in this study.**

**Table S5 Mutations identified in KomABC-resistant Rao1 mutants**

**Table S6 Proteins identified by the mass spectrometry of the samples co-purified with His-tagged KomC and untagged KomC.**

**Table S7 Peptides and proteins identified by mass spectrometry analysis of the gel slice in Fig. 2C.**
